# Spatio-temporal dynamics of lateral Na^+^ diffusion in apical dendrites of mouse CA1 pyramidal neurons

**DOI:** 10.1101/2025.08.06.668873

**Authors:** Joel S. E. Nelson, Jan Meyer, Niklas J. Gerkau, Karl W. Kafitz, Ghanim Ullah, Fidel Santamaria, Christine R. Rose

## Abstract

Sodium ions (Na^+^) are major charge carriers mediating neuronal excitation and play a fundamental role in brain physiology. Glutamatergic synaptic activity is accompanied by large transient Na^+^ increases, but the spatio-temporal dynamics of Na^+^ signals and properties of Na^+^ diffusion within dendrites are largely unknown. To address these questions, we employed multi-photon Na^+^ imaging combined with whole-cell patch-clamp in dendrites of CA1 pyramidal neurons in tissue slices from mice of both sexes. Fluorescence lifetime microscopy revealed a dendritic baseline Na^+^ concentration of ~10 mM. Using intensity-based line-scan imaging we found that local, glutamate-evoked Na^+^ signals spread rapidly within dendrites, with peak amplitudes decreasing and latencies increasing with increasing distance from the site of stimulation. Spread of Na^+^ along dendrites was independent of dendrite diameter, order or overall spine density in the ranges measured. Our experiments also show that dendritic Na^+^ readily invades spines and suggest that spine necks may represent a partial diffusion barrier. Experimental data were well reproduced by mathematical simulations assuming normal diffusion with a diffusion coefficient of 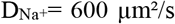. Modeling moreover revealed that lateral diffusion is key for the clearance of local Na^+^ increases at early time points, whereas when diffusional gradients are diminished, Na^+^/K^+^-ATPase becomes more relevant. Taken together, our study thus demonstrates that Na^+^ influx causes rapid lateral diffusion of Na^+^ within spiny dendrites. This results in an efficient redistribution and fast recovery from local Na^+^ transients which is mainly governed by concentration differences.

**Significance statement:** Activity of excitatory glutamatergic synapses generates large Na^+^ transients in postsynaptic cells. Na^+^ influx is a main driver of energy consumption and modulates cellular properties by modulating Na^+^-dependent transporters. Knowing the spatio-temporal dynamics of dendritic Na^+^ signals is thus critical for understanding neuronal function. To study propagation of Na^+^ signals within spiny dendrites, we performed fast Na^+^ imaging combined with mathematical simulations. Our data shows that normal diffusion, based on a diffusion coefficient of 600 µm^2^/s, is crucial for fast clearance of local Na^+^ transients in dendrites, whereas Na^+^ export by the Na^+^/K^+^-ATPase becomes more relevant at later time points. This fast diffusive spread of Na^+^ will reduce the local metabolic burden imposed by synaptic Na^+^ influx.

## Introduction

Sodium ions (Na^+^) are major charge carriers for the generation of action potentials and excitatory postsynaptic currents in the brain. Contrary to the long-held view that electrical signaling does not result in changes in the intracellular Na^+^ concentration ([Na^+^]_i_), it is now widely established that action potentials cause a transient [Na^+^]_i_ increase in axons and presynaptic terminals of central neurons (Lasser-Ross and Ross, 1992; Kole et al., 2008; Fleidervish et al., 2010; Zhu et al., 2020; Zang and Marder, 2021; Kotler et al., 2023). Activity-induced Na^+^ transients can also arise in dendrites and spines upon opening of voltage-gated Na^+^ channels during back-propagating action potentials, as e.g., shown for hippocampal CA1 pyramidal neurons (Jaffe et al., 1992; Rose et al., 1999). Particularly prominent Na^+^ transients are evoked by excitatory synaptic activity and Na^+^ influx through ionotropic glutamate receptors into postsynaptic dendrites and spines (Rose and Konnerth, 2001; Miyazaki and Ross, 2017; Gerkau et al., 2019; Miyazaki and Ross, 2022).

Intracellular Na^+^ signals have fundamental physiological consequences for neurons. Na^+^ is extruded by the Na^+^/K^+^-ATPase (NKA) against a large inward electrochemical gradient, which represents a major metabolic challenge for the brain (Sweadner, 1995). Earlier estimates revealed that Na^+^-based action potentials consume about 30%, whereas Na^+^ influx through ligand-gated channels requires up to 50% of cellular ATP production (Lennie, 2003; Hallermann et al., 2012; Harris et al., 2012; Lezmy et al., 2021). Furthermore, Na^+^ influx can result in a reduction of the driving force for Na^+^ inward currents (Sejnowski and Qian, 1992), and computational approaches suggested that [Na^+^]_i_ elevations might contribute to the termination of seizures (Krishnan and Bazhenov, 2011; Gonzalez et al., 2024). Increases in [Na^+^]_i_ also reduce the driving force for Na^+^-dependent secondary transporters and thereby decrease the export of protons through Na^+^/H^+^-exchangers, fostering cellular acidification(Zhao et al., 2016). Other transporters, like the Na^+^/Ca^2+^-exchanger (NCX) (Blaustein and Lederer, 1999) can even reverse in response to an elevation in [Na^+^]_i_. Such Na^+^-driven NCX reversal contributes to intracellular Ca^2+^ signalling, a phenomenon described for neuronal compartments as well as for astrocytes (Bouron and Reuter, 1996; Regehr, 1997; Scheuss et al., 2006; Song et al., 2013; Zylbertal et al., 2015; Ziemens et al., 2019; Rose et al., 2020).

Despite the high functional relevance of changes in [Na^+^]_i_ for neurons, knowledge about their spatio-temporal properties is extremely limited. It is for example known that recovery from global Na^+^ transients is mainly achieved by the NKA. There is also evidence that local Na^+^ transients, e.g., those accompanying synaptic activity and opening of ionotropic receptors in spiny dendrites and spines, are mainly cleared by fast lateral diffusion (Mondragao et al., 2016; Miyazaki and Ross, 2017). Moreover, earlier work has revealed high diffusion coefficients for Na^+^ in muscle cells or in large-caliber lizard axons 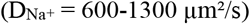 (Kushmerick and Podolsky, 1969; David et al., 1997). Diffusional properties of Na^+^ along dendrites, however, are practically unknown. Finally, a recent computational study has indicated that diffusion of Cl^-^ along dendrites is slowed down by the presence of dendritic spines (Mohapatra et al., 2016), but again, information on the relevance of spine density on Na^+^ diffusion in dendrites is lacking.

Here, we addressed these questions using multi-photon imaging in the line-scan mode in apical dendrites of CA1 pyramidal neurons in organotypic tissue slices of the mouse hippocampus. Brief iontophoretic application of glutamate served to induce a local increase in dendritic [Na^+^]_i_. Numerical simulations and mathematical modelling were employed to test the plausibility of our experimental findings and to reveal the biophysical properties of Na^+^ diffusion within dendrites. Our experiments and simulations show that the spread of Na^+^ in dendrites upon induction of local influx is governed by normal diffusion towards regions of lower Na^+^ concentrations at a 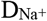 of 600 μm^2^/s. Moreover, we found that Na^+^ diffusion is primarily independent on the dendrite order or on the presence of spines.

## Material and Methods

### Ethical approval

The study was conducted in accordance with the guidelines of the Heinrich Heine University Düsseldorf, as well as the European Community Council Directive (2010/63/EU). All experiments were communicated and approved by the Animal Welfare Office at the Animal Care and Use Facility of the Heinrich Heine University (Institutional Act No. O52/05). In accordance with the recommendations of the European Commission (published in: Euthanasia of experimental animals, Luxembourg: Office for Official Publications of the European Communities, 1997; ISBN 92–827–9694-9), animals, all younger than 10 days, were killed by decapitation.

### Preparation of organotypic tissue slice cultures

Organotypic tissue slice cultures were prepared from Balb/C mice (postnatal days (P) 6-9; both sexes) following a modified protocol originally described by Stoppini and colleagues (Stoppini et al., 1991; Lerchundi et al., 2019). Tissue slice cultures for experiments illustrated in Fig. 1E (5 mM and 17 mM internal Na^+^) were prepared from C57BJ/6J mice. Animals were decapitated, and their brains quickly removed and placed into ice-cold artificial cerebral spinal fluid (ACSF) containing (in mM): 130 NaCl, 2.5 KCl, 2 CaCl_2_, 1 MgCl_2_, 1.25 NaH_2_PO_4_, 26 NaHCO_3_, and 10 glucose, bubbled with 95% O_2_/5% CO_2_, pH 7.4, and an osmolarity of 310 ± 5 mOsm/l. Subsequently, brains were separated into hemispheres and cut into 250 µm thick slices in a parasagittal orientation using a vibrating blade microtome (HM 650V, Thermo Fisher Scientific, Waltham, MA, USA).

**Figure 1:**
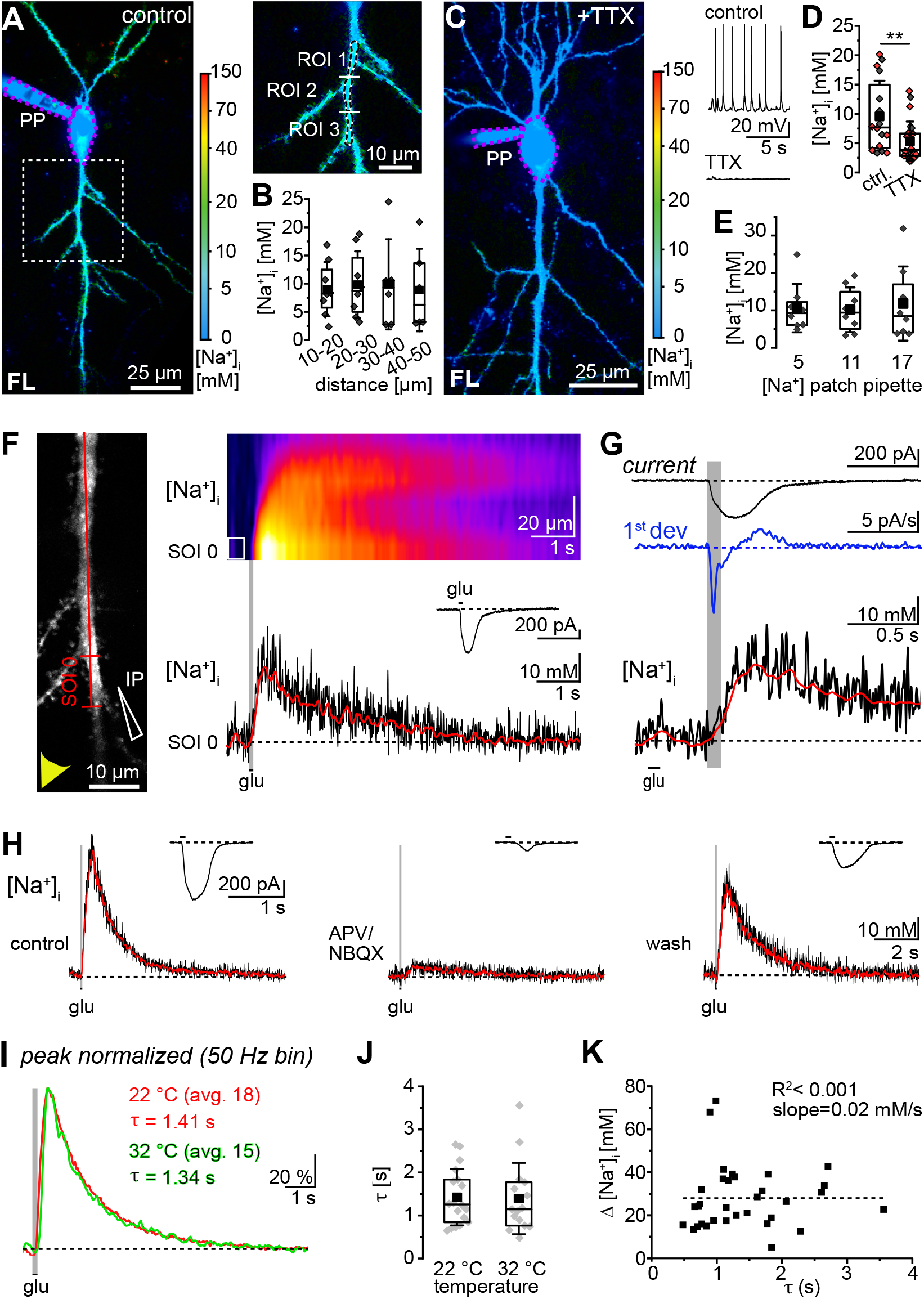
FLIM-based analysis of [Na^+^]_i_ and line scan recording of [Na^+^]_i_ changes. A: Color-coded image of ING2 fast Fluorescence Lifetime (FL) of a CA1 neuron. Color-code on the right illustrates [Na^+^]_i_ as determined by *in situ* calibration. The dotted box: region enlarged on the right; PP: patch pipette. Upper Right: dotted lines indicate 10 µm long ROIs from which FL was analyzed. Note that pipette and soma (encircled by magenta) are depicted with the same color code as dendrites, calibration properties, however, differ in these compartments. B: Box plots illustrating baseline [Na^+^]_i_ along primary dendrites at different distances from the soma (n=8, N=6). C: Color-coded image of ING2 FL of a cell exposed to TTX. Again, pipette and soma are depicted with the same color code as dendrites, but calibration properties differ. Upper Right: current-clamp recording in control and with TTX. D: Box plots showing [Na^+^]_i_ in control (16 recordings in 8 cells) and with TTX (24 recordings in 12 cells; p=0.006, Mann-Whitney test). E: Box plots showing [Na^+^]_i_ in primary dendrites in control conditions from cells patched with a pipette saline containing either 5, 11 or 17 mM Na^+^. B, D and E: Shown are single data points (diamonds), mean (squares), median (horizontal line), quantiles (box), and SD (whiskers). D: Grey diamonds: primary dendrites, red diamonds: secondary dendrites; statistical significance in indicated by asterisks with 0.01<**p<0.001. F: Left: Projection of an SBFI-filled primary dendrite. Red line: position of the scan line; SOI 0: segment exhibiting the largest relative change in fluorescence. IP: iontophoresis pipette; arrowhead indicates flow of bath perfusion. Top Right: False color-coded (x,t) image of the baseline-corrected line scan, depicting [Na^+^]_i_ changes induced by glutamate iontophoresis (100 ms). Vertical box: position of SOI 0. Center: Glutamate-induced somatic inward current. Bottom Right: Glutamate-induced change in [Na^+^]_i_ in SOI 0. Black: baseline corrected trace (subjected to 50 Hz low pass FFT), red: filtered trace as given by the Python-based program. G: Somatic inward current, its calculated first deviation and the accompanying change in [Na^+^]_i_ in SOI 0. H: Glutamate-induced dendritic [Na^+^]_i_ transients and somatic currents taken under control conditions (left), after wash in of glutamate receptor blockers APV and NBQX (middle), and after washout of the blockers (right). I: Peak-normalized, averaged traces from experiments conducted at 22 °C (n=18, N=18; red) and at 32 °C (n=15, N=11; green). J: Boxplot comparing the decay time constants (τ) of [Na^+^]i recovery at 22 °C and 32 °C. Shown are individual data points (diamonds), means (squares), medians (horizontal lines), quantiles (boxes), and standard deviations (whiskers). Data was statistically analyzed using a Mann-Whitney test (p=0.704). K: Scatterplot showing correlation between Ω[Na^+^]_i_ and τ of the signal measured at SOI 0 in experiments taken from DIV 10-25 primary dendrites. The dotted line shows a linear fit (R^2^ < 0.001; slope=0.02).

The hippocampus and part of the adjacent cortex were isolated, washed in Hanks’ balanced salt solution (Sigma-Aldrich, St. Louis, Mo, USA) and transferred onto Biopore membranes (Millicell standing insert, Merck Millipore, Darmstadt, Germany). Membranes were then placed into a 6 well plate and maintained in an incubator at the interface between humidified air containing 5% CO_2_ and culture medium at 36°C. Culture medium contained the following: minimum essential medium (MEM; Sigma-Aldrich), 20% heat-inactivated horse serum (Thermo Fisher Scientific), 1 mM L-glutamine, 0.01 mg/ml insulin, 14.5 mM NaCl, 2 mM MgSO_4_, 1.44 mM CaCl_2_, 0.00125% ascorbic acid, and 13 mM D-glucose. Slices were kept in the incubator until use between 3-50 days *in vitro* and the medium was changed every 3 days. Unless otherwise mentioned, experiments were carried out at room temperature (22 ± 1°C), and slices were constantly perfused with standard ACSF at a rate of 2.5 ml/min. In experiments conducted at 32 ± 1°C, slices were perfused with ACSF containing (in mM): 138 NaCl, 2.5 KCl, 2 CaCl_2_, 1 MgCl_2_, 1.25 NaH_2_PO_4_, 18 NaHCO_3_ and 10 glucose, bubbled with 95% O_2_/5% CO_2_, pH 7.4.

### Electrophysiology, dye loading and glutamate iontophoresis

Whole-cell patch-clamp measurements were performed using a pipette solution containing (in mM): 114 KMeSO_3_, 32 KCl, 10 HEPES (4-(2-hydroxyethyl)-1-piperazineethanesulfonic acid), 10 NaCl, 4 Mg-ATP, 0.4 Na_3_-GTP; pH adjusted to pH 7.3. To study the influence of pipette [Na^+^] on dendritic [Na^+^]_i_, pipette solutions contained either 4 or 16 mM NaCl with KCl concentrations adapted to maintain isoosmolarity. Cells were held in the current-clamp mode in “zero current” conditions or in the voltage-clamp mode at −70 mV (liquid junction potential not corrected) using an EPC10 amplifier (HEKA Elektronik, Lambrecht, Germany) and PatchMaster-Software (HEKA Elektronik). Electrophysiological data was analyzed using OriginPro software (OriginPro 2021, Northampton, MA, USA).

For multiphoton imaging of Na^+^, the membrane-impermeable forms of the fluorescent chemical Na^+^ indicators ION-NaTRIUM-Green-2 (ING2 TMA^+^ salt; IonBiosciences/Mobitec, Göttingen, Germany) or Na^+^-binding benzofuran-isophthalate (SBFI K^+^ salt; Teflabs, Austin, TX, USA) were added to the intracellular saline at a final concentration of 50 µM (ING2) or 1 mM (SBFI). The dyes were loaded into individual neurons using whole-cell patch-clamp for at least 30 minutes before starting imaging experiments as reported previously (Gerkau et al., 2019). Standard visualization of filled neurons was performed using maximum intensity projections from z-stacks (1 µm steps).

For three-dimensional reconstruction of dye-filled dendrites, z-stacks were taken at 0.2 µm spacing between focal planes. Z-stacks were reconstructed and visualized using the ImageJ software (National Institutes of Health, USA). Stacks were imported into the Huygens Software (Huygens Professional, SVI imaging, Hilversum, Netherlands) for deconvolution using an experimentally determined point spread function (PSF). Deconvoluted stacks were reconstructed and projected as a three-dimensional structure with the Imaris software (Center for Bio-Image Informatics, University of California, Santa Barbara, USA). Dendrites were traced and reconstructed using the Imaris filament tool to determine their diameter and to identify presumptive spines.

For local iontophoresis of glutamate, a fine-tipped, high resistance (90-120 MΩ) borosilicate glass pipette was pulled out (PP-830, Narishige; Tokyo, Japan) and filled with 150 mM glutamate. Pipettes were coupled to an iontophoresis module (MVCS-M-45, NPI Electronic GmbH, Tamm, Germany) and a retain current was applied to prevent leakage of glutamate. The pipette tip was lowered into the tissue slice under visual control and placed close to a chosen dendrite. The tip of the iontophoresis pipette was positioned upstream relative to the bath perfusion and directed against the perfusion flow to minimize the extracellular diffusion of glutamate along the chosen dendritic section.

### Fluorescence lifetime imaging of Na^+^ with ING2

Fluorescence lifetime imaging (FLIM) of ING2 was performed as described earlier (Meyer et al., 2022), employing a laser-scanning microscope based on a Fluoview300 system (Olympus Deutschland GmbH, Hamburg, Germany), or a Nikon A1-R MP system (Nikon Europe, Amsterdam, The Netherlands), equipped with a water immersion objective (NIR Apo 60x, NA 1.0, Nikon Instruments Europe, Düsseldorf, Germany). Laser pulses (100 fs) were generated at 80 MHz by a mode-locked Titan Sapphire laser (Mai Tai or Mai Tai DeepSee, Newport, Spectra Physics; Irvine, CA, USA); excitation wavelength was 840 nm. Amplitude weighted, average fluorescence lifetimes (FL) were measured using time-correlated single photon counting (TCSPC) with a spatial resolution of 0.57 × 0.57 µm per pixel and analyzed employing MultiHarp 150 electronics and SymPhoTime 64 acquisition software version 2.8 (both PicoQuant GmbH; Berlin, Germany). Fluorescence emission was bandpass-filtered (534/30 or 540/25 nm, AHF Analysentechnik; Tübingen, Germany) and directed to a phosphomolybdic acid (PMA) hybrid photodetector (PicoQuant GmbH; Berlin, Germany). Images of dye-filled neurons represent fast lifetime (fLT) images as generated by the SymPhoTime64 software (Meyer et al., 2022). Data analysis was performed using average fluorescence lifetimes (τ_AVG_) from defined regions of interest (ROIs) drawn along dendrites. Decay curves were analyzed by reconvolution of the instrument response function (IRF) and bi-exponential fitting resulting in the amplitude weighted τ_AVG_ (Meyer et al., 2022).

For determination of dendritic Na^+^ concentrations ([Na^+^]_i_), z-stacks (1 µm steps, each 15 or 30 frames) were taken. Photons were collected until >1,000,000 photons were reached for each plane, and maximal projections of fLT images were calculated. Further analysis of the extracted τ_AVG_, including conversion of FL into [Na^+^]_i_ (see below) and statistical analysis of the data, was commenced using Excel (Excel 2019, Microsoft Corporation, Redmond, WA, USA) and Origin software (OriginPro 2021, Northampton, MA, USA).

To convert FL into [Na^+^]_i_, ING2 was calibrated *in situ* as described earlier (Meier et al., 2006; Langer and Rose, 2009; Meyer et al., 2022). The properties of Na^+^ indicator dyes (including apparent binding affinity to Na^+^) are different under *in vitro* conditions as compared to those inside cells (Despa et al., 2000; Langer and Rose, 2009; Lamy and Chatton, 2011; Naumann et al., 2018; Meyer et al., 2019). Indeed, when employing *in situ* calibration parameters determined for ING2, the FL obtained from ROIs positioned over patch pipettes were too low for a conversion into meaningful [Na^+^]. In contrast, using *in vitro* calibrations parameters established earlier (Meyer et al., 2022) resulted in a value of 11.0 mM that was in good agreement with the known pipette [Na^+^] (11.2 mM). In somata, *in vitro* calibration parameters resulted in rather high apparent [Na^+^]_i_ (> 40 mM), while *in situ* calibration indicated an apparent [Na^+^]_i_ significantly below that of the pipette solution (0-5 mM). These results suggest that in cell bodies, held in whole-cell patch-clamp mode and thus in direct contact with the pipette saline, neither of the two calibrations is suited to reliably convert FL into [Na^+^]_i_. Our FL analysis, therefore, did not consider somata.

### Fluorescence intensity imaging of SBFI and automated analysis of line scans

Imaging was performed at the multi-photon system described above for FLIM at an excitation wavelength of 790 nm. Fluorescence emission <700 nm was directed to the internal Fluoview photomultiplier tubes (Olympus, Hamburg, Germany). Line scans were performed on dye-filled dendrites with a scanning frequency of 233-747 Hz. ImageJ was used to generate line scan projections and projections of z-stacks which were later used for 3-dimensional morphological analysis.

Unless stated otherwise, line scans were analyzed using a custom Python-based script for unbiased detection of peak time point and peak amplitude of fluorescence changes (code see Repositorium/Github Link). The program was operated through the Spyder software (version 5.2.2, Austin TX, USA) over anaconda navigator (version 2.3.2, Austin TX, USA). For this, line scan images were converted into text files using ImageJ and Excel software (Microsoft Corporation, Redmond, WA, USA), resulting in a pixel-specific listing of fluorescent intensity values over time. The files were then analyzed by the script.

First, the program established segments of interest (SOIs) by averaging intensity values of *n* neighboring pixels along the line scan, e.g., for an *n* of 50, starting from pixel 1 to 50, 51 to 100, and so forth. Next, the average baseline fluorescence intensity (F_0_) of each SOI was determined from the first 500 ms of each recording (during which no stimulation was performed), after which traces were normalized to F_0_ (ΔF/F_0_). Subsequently, an automated correction for photo-bleaching was performed by fitting a bi-exponential decay function 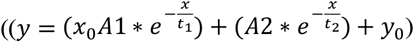 with t_1_ >0.2 and t_1_ + t_2_ < 10) through the data points between 0-500 ms (“baseline period”) and the last quarter of each trace (i.e. when changes in fluorescence induced by glutamate had fully recovered) and subtracting this fit from the respective data trace. Afterwards, traces were filtered using a Butterworth low pass filter (cutoff: 1.5; sampling frequency: 40 Hz; order: 3) and a rolling average (window of 25 data points).

Finally, the peak time point and peak amplitude of glutamate-induced changes in fluorescence intensity were determined by automatically extracting the lowest point of each trace (note that SBFI intensity decreases with increasing [Na^+^]). This point had to obey the following criteria: 1) signal-to-noise ratio (calculated by the division of the peak by the 1% percentile of the trace) larger than 20%; 2) occurrence between 500 ms (time point at which iontophoresis was triggered) and 3000 ms; 3) baseline drift less than 10%; 4) peak amplitude larger than two times the standard deviation of the noise during baseline conditions. If these criteria were not met, the data trace was discarded. The remaining traces (unfiltered baseline-corrected traces, Butterworth-filtered traces and traces subjected to the rolling average filter) were then plotted using Origin software (OriginPro 2021). Moreover, peak amplitudes and time points determined by the program were imported into the Origin software for further analysis.

### Analysis of Na^+^ diffusion within dendrites

Lateral Na^+^ diffusion in dendrites was analyzed following procedures similar to those described by Santamaria et al. (Santamaria et al., 2006). First, we established SOIs of 1.02 µm length (5 pixels) along dendritic line scans using the Python-based automated analysis and performed a temporal binning to obtain frequencies of 50 Hz. Furthermore, conversion of ΩF/F_0_ values into changes in [Na^+^]_i_ was performed via the software, using calibration parameters established as described above and assuming the baseline [Na^+^]_i_ as determined before by FLIM (see above and results). As a last step, measurements were normalized using Eq. 1:

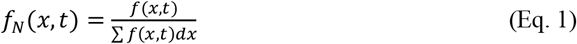

where *f*_*N*_*(x,t)* is the normalized concentration distribution over the length of the imaged dendrite. Concentration distributions (*f(x,t)*) were thereby taken every 0.02 s (*dt* = 0.02 s) and each SOI had a size of *dx* = 1.02 µm (5 pixels). Measurements from a given dendrite order (primary or secondary dendrite) and time in culture were then pooled for further analysis.

For calculation of diffusion coefficients we assumed a one-dimensional diffusion process along the axis of the dendrite (Santamaria et al., 2006), which is described by Eq. 2:

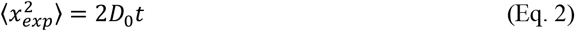

where ⟨*x*^*2*^_*exp*_⟩ is the variance at each time point, *D*_*0*_ the diffusion coefficient, and *t* is time. The variance was calculated for each time point using Eq. 3:

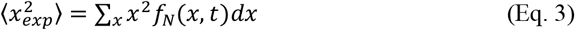

We implemented this numerical integration in Matlab using a trapezoidal integration method. Finally, the apparent diffusion coefficient *D*_*app*_ was calculated as

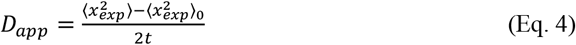

where ⟨*x*^*2*^_*exp*_⟩_*0*_ is the variance at the initial condition.

In our experiments, the increase in dendritic [Na^+^]_i_ was induced by iontophoretic glutamate application which did not represent a strictly localized point source for Na^+^.

Moreover, glutamate application resulted in a Na^+^ influx for several hundreds of ms. Resulting from that, the rising phase of the signal consists of both an influx and lateral diffusion of Na^+^. We assume that after the peak is reached, the spread of Na^+^ is due to diffusion. Therefore, the concentration distribution which was measured at the peak of the overall signal was defined as initial distribution.

To further verify our results and to assess the effect of boundary conditions set by the experiments such as the duration of Na^+^ influx and the restricted length of dendritic segments imaged, we simulated a normal 2D diffusion process using:

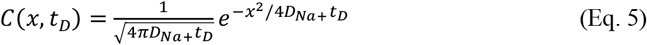

where *C(x,t*_*D*_) is the concentration spread over the length of the dendrite at a certain time (*t*_*D*_*), x* is distance and D_Na+_ is either 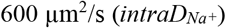, a value reported earlier for unhindered Na^+^ diffusion in cellular structures (Kushmerick and Podolsky, 1969) or was simulated for the most effective fit of the experimental data 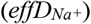. The variance ⟨*x*^*2*^⟩ is:

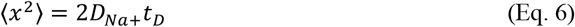

To take the initial distribution into account, we calculated the time which the simulation needed to reach that distribution (*t*_*shift*_) given ⟨*x*^*2*^_*exp*_⟩_*0*_:

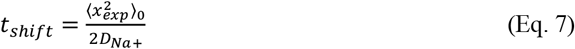

We then simulated a normal diffusion process for 3 s after *t*_*shift*_ using Eq. 5. Thereby, *t*_*shift*_ was associated to the peak timepoint (*t =0*) of the experiment. For calculation of the variance, performed using Eq. 6, we took the length of the measured dendrite into account. Results of the simulation thus show the variance which is expected within the boundary conditions of the experiment during a normal diffusive process with an 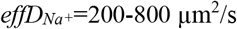 (Table 1).

**Table 1:**
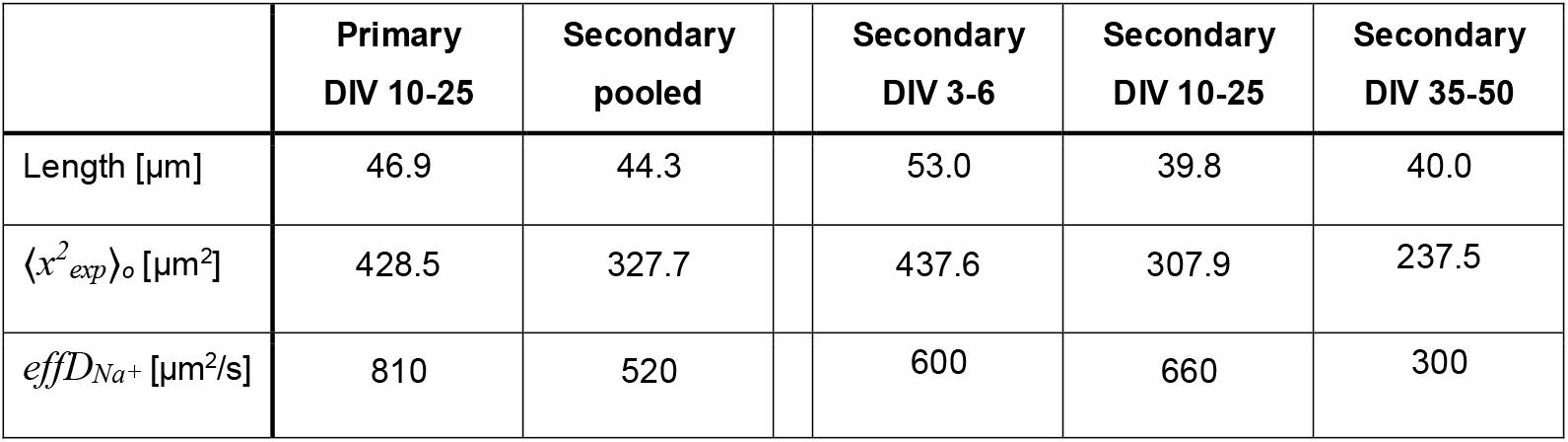
Initial variance of [Na^+^]_i_ and effective D^Na+^ for all conditions. Properties of the experiments incorporating the measured length and the determined initial variance (⟨*x*^*2*^_*exp*_⟩_*o*_) for every experimental condition. Lastly the listing of the determined effective 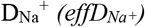 for each condition.

|  | Primary<br>DIV 10-25 | Secondary<br>pooled |  | Secondary<br>DIV 3-6 | Secondary<br>DIV 10-25 | Secondary<br>DIV 35-50 |
| --- | --- | --- | --- | --- | --- | --- |
| Length [$\mu m$] | 46.9 | 44.3 | | 53.0 | 39.8 | 40.0 |
| $\langle x^2_{exp} \rangle_o$ [$\mu m^2$] | 428.5 | 327.7 | | 437.6 | 307.9 | 237.5 |
| $effD_{Na^+}$ [$\mu m^2/s$] | 810 | 520 | | 600 | 660 | 300 |

### Simulation of lateral spread of Na^+^

To simulate the lateral spread of Na^+^ in dendrites, we used a reconstructed morphology of a pyramidal neuron, obtained from Neuromorpho.org (RRID:SCR_002145; (Buchin et al., 2022); morphology #NMO_276156) with a total of 2895 compartments (Tecuatl et al., 2024). The morphology file contains an integer as compartment identifier, type of compartment (soma, axon, basal dendrite, apical dendrite, etc.), coordinates (x, y, z), radius, and the parent compartment of each compartment. The coordinates of two adjacent compartments were used to calculate the length of each compartment. Thus, the length and radius vary from compartment to compartment according to the morphological reconstruction. The compartment’s identifier and parent compartment’s identifier were used to couple different compartments according to their positioning in the morphology. Overall, this information is enough to spatio-temporally model the spread of [Na^+^]_i_ using the actual morphology where the currents are distributed following the reported electrophysiology of pyramidal cells (Spruston, 2008). As explained below, although the effect of the radius of each compartment is incorporated in the electrical coupling between neighboring compartments, the diffusion of Na^+^ flux between neighboring compartments only depend on *intraD*_*Na+*_, separation between two compartments, and the concentration gradient between the neighboring compartments. The treatment of a compartment as a point follows from the assumption that Na^+^ stabilizes very quickly across the width of the compartment due to the large value of *intraD*_*Na+*_.

The neuron was modeled using Hodgkin-Huxley type conductance based current equations together with dynamic intra- and extracellular ion concentrations. The reversal potentials for various currents are made variable such that they depend on the instantaneous values of concentrations of relevant ions inside and outside the cell. The membrane potential of i^th^ compartment, *V*^*(i*^,^)^ is modeled with the following rate equation.

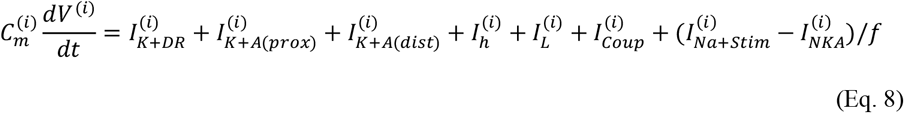

where the equations for the delayed rectified K^+^ (*I*_*K+DR*_), proximal A (*I*_*K+A(prox)*_), distal A (*I*_*K+A(dist)*_), and h (*I*_*h*_) currents are taken from (Migliore et al., 1999; Migliore et al., 2004) and are given as follows. We remark that ignoring *I*_*K+A(prox)*_, *I*_*K+A(dist)*_, and *I*_*h*_ currents does not change results from the simulation significantly and that these currents are included only for completion, considering the previously reported currents in different compartments (Spruston, 2008). The leak (*I*_*L*_) and coupling (*I*_*Coup*_) currents between neighboring compartments are also given below. *I*_*Na+Stim*_ and *I*_*NKA*_ represent localized Na^+^ stimulus (Na^+^ is injected to raise [Na^+^]_i_ locally) and NKA, respectively. *f* = 0.0445 (Barreto and Cressman, 2011) converts current from μA/cm^2^ to mM/sec. Note that our experiments are performed in the presence of TTX, therefore, fast voltage-gated Na^+^ current is not included in the model. Note that in the equations for different currents, we have dropped the superscript *(i)* for clarity, however, these currents are computed using voltage, intra- and extracellular ion concentrations of that compartment.

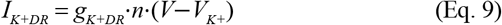

where *g* _*K+DR*_ = 40 mS/cm^2^.

The gating variable, *x (*e.g., *n, l*, etc.) for various currents are given by the following functional form:

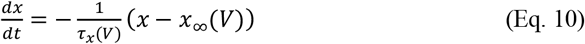

with *n*_∞_ = 1/(1 + *α*_*n*_); *τ* _*n*_ = 50*β*_*n /*_(1 + *α*_*n*_); *α*_*n*_ = exp(*−*11(*V −* 13)); *β*_*n*_ = exp(*−*08(*V −* 13)); *τ* _*n*_ = 2 ms if *τ* _*n*_ *<* 2 ms.

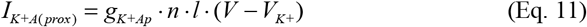

Where g_K+Ap_= 48 mS/cm^2^, 48 mS/cm^2^ and 4.8 mS/cm^2^ for soma, basal dendrites, and axon, respectively (note that the morphology file contains information about the type of compartment).

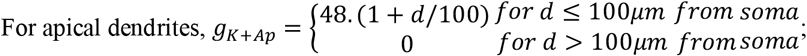

*n*_∞_ = 1/(1 + *α*_*n*_); *τ* _*n*_ = 4*β*_*n /*_(1 + *α*_*n*_); *α*_*n*_ = exp(*−*0.038(1.5 + 1 / (1 + exp(*V* + 40) / 5)) · (*V −* 11)); *β*_*n*_ = exp(*−*0.038(0.825 + 1 / (1 + exp(*V* + 40) / 5)) · (*V −* 11)); *l*_∞_ = 1/(1 + *α*_*l*_); *τ* _*l*_ = 0.26 · (*V* + 50); *α*_*l*_ = exp(0.11(*V* + 56)); *τ*_*n*_ = 1ms if *τ*_*m*_ *<* 1 ms; *τ* _*l*_ = 2 ms if *τ* _*l*_ *<* 2 ms. The distance (*d*) of a compartment from the soma is calculated using the (x, y, z) coordinates of both compartments.

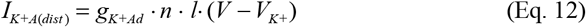

*I*_*K+A*(*dist*)_ ·is only considered in apical dendrites with:

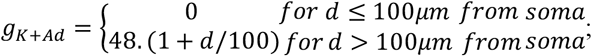

*n*_∞_ = 1/(1 + *α*_*n*_); *τ* _*n*_ = 2*β*_*n /*_(1 + *α*_*n*_); *α*_*n*_ = exp(*−*0.038(1.8 + 1 / (1 + exp(*V* + 40) / 5)) · (*V* + 1)); *β*_*n*_ = exp(*−*0.038(0.7 + 1 / (1 + exp(*V* + 40) / 5)) · (*V* + 1)); *l*_∞_ = 1/(1 + *α*_*l*_); *τ* _*l*_ = 0.26 · (*V* + 50); *α*_*l*_ = exp(0.11(*V* + 56)); *τ*_*n*_ = 1*ms* if *τ*_*m*_ *<* 1 ms; *τ* _*l*_ = 2 ms if *τ* _*l*_ *<* 2 ms.

I_h_ is included in the soma and apical dendrites, and is given by

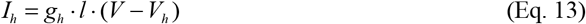

where *V*_*h*_ *=* −30mV and *g*_*h*_· = 0.05 · (1 + 3*d* / 100). Here *d* is the distance from the soma.

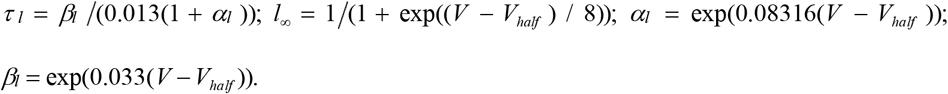

The leak current, *I*_*L*_ is composed of three components; Na^+^ leak (*I*_*L,Na+*_), K^+^ leak (*I*_*L,K+*_), and Cl^-^ leak (*I*_*L,Cl-*_).

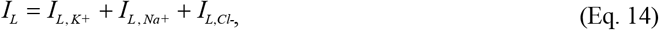

And *I*_*L, K+*_ = *g*_*L*_ (*V − V*_*K+*_); *I*_*L,Na+*_ *= 0*.*35g*_*L*_ *(V – V*_*Na+*_*); I*_*L,Cl-*_*= g*_*L*_ *(V – V*_*Cl-*_); where *g*_*L*_ = 10^3^/R_m_ is the leak conductance and *R*_*m*_ = 50,000 Ωcm^2^ is the resistivity of the membrane.

Neighboring compartments are electrically coupled with each other through a coupling current (*I*_*Coup*_).

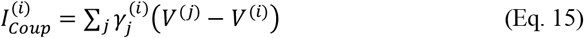

where the summation is over the nearest neighbors. The identifiers of a compartment and its parent compartment are used to calculate the number of nearest neighbors. If compartment (*i*) has length *L*^*(i)*^ and radius *r*^*(i)*^ and compartment (*j*) has length *L*^*(j)*^ and radius *r*^*(j)*^ then the intercompartmental resistance is the sum of the two resistances from the middle of each compartment with respect to the junction between them, i.e. 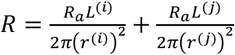, where *R*_*a*_ = 400 Ωcm^2^ is the intracellular resistivity. To compute intercompartmental conductance *γ* we take the inverse of the above expression and divide the result by the total area of compartment (*i*) (Dayan and Abbott, 2001).

The dynamics of intracellular Na^+^ in the i^th^ compartment and K^+^ in its corresponding extracellular compartment (*K*_*o*_) are formulated by the following equations (Cressman et al., 2009).

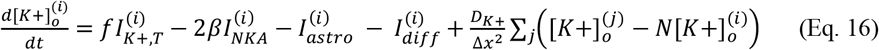

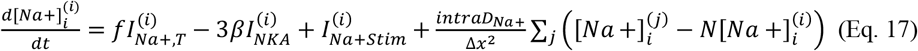

As mentioned above, *f* converts current density into flux. *I*_*Na+,T*_ and *I*_*K+,T*_, respectively are the total Na^+^ and K^+^ currents in the compartment or its corresponding extracellular compartment, due to different channels described above. That is,

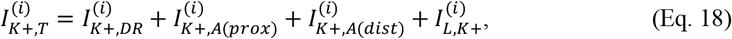

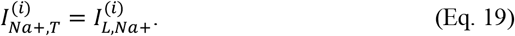

We consider the ratio of intra-to the extracellular volume for each compartment (*β*) to be 6. *I*_*NKA*_, *I*_*astro*_, and *I*_*diff*_ represent NKA, buffering of K^+^ by an astrocyte from the extracellular space of the compartment, and diffusion of K^+^ between the extracellular space of the compartment and bath solution. In other words, each intracellular compartment has its corresponding extracellular compartment, which has its associated astrocytic compartment and diffusion flux with the bath solution. As before, although we have dropped the superscript *(i)* for clarity, ion fluxes for a given compartment are calculated using the ion concentrations in that compartment and its corresponding extracellular space. *I*_*Na+Stim*_ represents intracellular injection of Na^+^ in a ~5 μm segment of the dendrite for 500 ms to raise local [Na^+^]_i_ to the desired value.

The equations for NKA and K^+^ diffusion between the extracellular space and bath are given by (Cressman et al., 2009; Ullah et al., 2009).

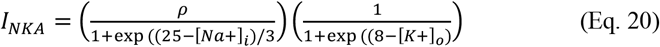

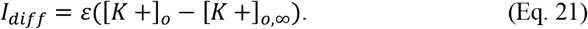

The K^+^ uptake by astrocytes is modeled by the following equation.

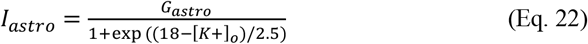

where *ρ* = 1.25 mM/s and *G*_*astro*_ = 67 mM/s is the maximum strength of NKA and astrocytic buffering, respectively. *ε* = 1.33/*s* is the diffusion constant for K^+^ from the extracellular space to the bath solution and [*K+*]_*o*,∞_ is the steady state K^+^ concentration in the bath and is set to 3 mM under physiological conditions.

The last terms in equation (16) and (17) represent the diffusion of extracellular K^+^ and intracellular Na^+^ between the neighboring compartments with diffusion coefficient of *D*_*K+*_ = 250μm^2^/s and *intraD*_*Na+*_ = 600 μm^2^/s, respectively. Δ*x* is the separation between two neighboring compartments, calculated using their (x, y, z) coordinates. No-flux boundary conditions are applied to the ending compartments. For example, a compartment at the end of a branch can exchange Na^+^ with its parent compartment (since the compartments at branch ends do not have daughter compartments). The same approach is also applied to the extracellular compartments. Furthermore, the movement of Na^+^ due to electric drift (electromigration) is ignored as it is significantly smaller than the Fickian diffusion and does not affect our results qualitatively (not shown).

Following (Cressman et al., 2009), we assume that the net flow of Na^+^ into the cell is matched by the flow of K^+^ out of the cell, thus giving the intracellular K^+^ concentration [K^+^]_i_:

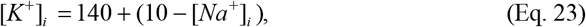

where the constants 140 mM and 10 mM represent the values for the intracellular K^+^ and Na^+^ concentrations at rest. We also assume that the total amount of Na^+^ is conserved:

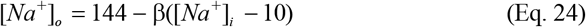

where [*Na*^*+*^]_*o*_ = 144 *− β*([*Na*^*+*^]_*i*_ *−* 10) (Eq. 24) where [Na^+^]_o_ represents the instantaneous value of extracellular Na^+^ and 144 mM is the extracellular Na^+^ concentration at rest. We remark that our model is a very simple representation of a complex reality. In addition to the simplification employed in Eqs. 23 and 24 and ignoring Cl^-^ dynamics and the effect of electric drift on the diffusion, we lumped K^+^ exchange between the ECS and astrocytes into a simple sigmoidal function, ignoring the complex behavior of various channels and other ion species. However, adding such complexity to the model would not change our main conclusions about the diffusion of intracellular Na^+^ in the neuron. Furthermore, we have shown previously that the intra- and extracellular ion dynamics from this simpler formalism closely matches that from the more extended model where [*Na*^*+*^]_*o*_, [*K*^*+*^]_*i*_, and Cl^-^ are modeled explicitly (Wei et al., 2014).

The reversal potentials for different ion currents are determined by the instantaneous values of their intra- and the extracellular concentrations according to the Nernst equation, with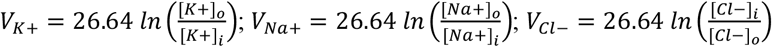.

The above equations were solved in Matlab version 2022b using Euler method with a timestep of 0.002 ms and were also confirmed using Python. Both versions of the code are available from the authors upon request.

### Experimental design and statistical analysis

Experiments were performed on tissue derived from neonatal animals of both sexes. Each set of experiments was performed on at least 4 different slices obtained from at least three different animals. Unless specified differently, *n* represents the number of analyzed cells; *N* represents the number of slices. Power analysis was conducted using G*Power 3.1.9.7. The effect size was calculated as the difference of means divided by the standard deviation of the control group. The α error probability was set to <0.05 with a resulting minimum power of 0.8.

Statistical analysis was performed by the OriginPro software (OriginPro 2021). Datasets were first tested for normal distribution using the Shapiro Wilk test. Normally distributed data sets were statistically analyzed by one-way ANOVA followed by a post hoc Bonferroni for unpaired datasets and a paired students t-test for paired data sets. Unpaired data which was not normally distributed was analyzed with a Mann-Whitney (MWU) test. Data are illustrated in box-and-whisker plots or scatter (x,y) plots unless otherwise specified. Box-and-whisker plots indicate the median (line), mean (square), quantile range (25/75, box) and SD (whiskers); p-values are indicated as follows: *: p< 0.05; **: 0.05> p >0.01 and ***: p< 0.001.

## Results

### Determination of dendritic [Na^+^]_i_ by FLIM

Former studies reported a baseline [Na^+^]_i_ of 10-15 mM in somata of CA1 pyramidal neurons. (e. g. (Langer and Rose, 2009; Kelly and Rose, 2010; Mondragao et al., 2016; Meyer et al., 2022). Information on dendritic [Na^+^]_i_, however, is still lacking. As local differences in [Na^+^]_i_ will influence its diffusion, we first employed FLIM of ING2 (Meyer et al., 2022) to obtain a direct, quantitative measure of [Na^+^]_i_ in apical dendrites. To this end, neurons were subjected to whole-cell patch clamp, revealing a resting membrane potential of −70 mV ± 3 mV (median=-70 mV; n=19, N=19). After cells were loaded with the dye, z-stacks encompassing soma and proximal apical dendritic tree were recorded using FLIM (Fig. 1A, C).

[Na^+^]_i_ was determined in 10 µm long ROIs drawn around primary dendrites, starting at 10 µm distance from the soma, as well as from secondary dendrites emerging from primary dendrites (Fig. 1A,B). We found no differences between ROIs at different distances from the soma, nor between different dendrite orders (Fig. 1A, B; Table 2). Data were thus pooled, revealing an average dendritic [Na^+^]_i_ of 9.6 ± 6.1 mM (median=7.7 mM; 8 measurements from primary and secondary dendrites each; n=8, N=6) (Table 2; Fig. 1D). Perfusion of slices with tetrodotoxin (TTX, 0.5 µM), suppressed spontaneous action potential firing (Fig. 1C), while resting membrane potentials remained unaltered (−70 mV ± 5 mV; median=-69 mV; n=19, N=19; p=0.738). In the presence of TTX, dendritic [Na^+^]_i_ was significantly lower than in control, averaging 5.3 ± 3.4 mM (median=3.9 mM; 24 ROIs pooled from primary and secondary dendrites; n=12, N=8; p=0.007) (Fig. 1C,D). The latter results also indicated that dendritic [Na^+^]_i_ was not governed by the pipette saline, containing 11 mM (11.2 mM, see methods) [Na^+^].

**Table 2:** Baseline [Na^+^]_i_ in primary and secondary apical dendrites. [Na^+^]_i_ (in mM), determined in 10 µm long ROIs along dendrites encompassing a distance of 10-50 µm from the soma (primary dendrites) or from the branching point from the primary dendrite, respectively. For statistical analysis between neighboring regions, normal distribution of the data was determined by the Shapiro Wilk test and one-way ANOVA with Bonferroni *post hoc* analysis was used for normal distributed data. Otherwise, Mann-Whitney tests were used. *P*-values are enclosed. For averages, ROIs from the entire length of the dendrite of one cell were pooled, before averaging the data. With n= number of ROIs and N=number of cells.

| Dendrite order | $[Na^+]_i$ [mM] | | | | | |
| --- | --- | --- | --- | --- | --- | --- |
| | Distance from soma (μm) | | | | Pooled [mM] | |
|  | 10-20 | 20-30 | 30-40 | 40-50 |  |  |
| Primary | 9.0 ± 4.8<br>(n=8; N=8) | 9.8 ± 5.9<br>(n=8; N=8;<br>p=0.77) | 9.9 ± 8.0<br>(n=6; N=6;<br>p=0.99) | 8.9 ± 7.3<br>(n=6; N=6;<br>p=0.82) | 10.2 ± 6.0 | p=0.7 |
| Secondary | 10.0 ± 8.4<br>(n=32; N=8) | 9.4 ± 6.2<br>(n=26; N=8;<br>p=0.96) | 9.6 ± 7.7<br>(n=14; N=8;<br>p=0.71) | 6.9 ± 4.4<br>(n=11; N=7;<br>p=0.60) | 9.0 ± 6.5 | |
| Average | 9.8 ± 7.8<br>(n=40; N=8) | 9.5 ± 6.1<br>(n=34; N=8;<br>p=0.92) | 9.7 ± 7.5<br>(n=20; N=8;<br>p=0.71) | 7.6 ± 5.5<br>(n=17; N=7;<br>p=0.53) | 9.6 ± 6.1 | |

To further assess whether dendritic [Na^+^]_i_ was influenced by the [Na^+^] of the patch pipette, experiments were performed with pipette salines containing different internal [Na^+^]. For a pipette [Na^+^] of 5 mM (5.2 mM), dendritic [Na^+^]_i_ was 10.6 mM (median=9.3 mM; n=8; N=5). Cells patched with a pipette saline containing 17 mM [Na^+^] (17.2 mM), had an average dendritic [Na^+^]_i_ of 11.8 mM (median=8.4 mM; n=8; N=4; p=0.793) (Fig. 1E). These results clearly demonstrate that [Na^+^]_i_ in dendrites was not clamped by the [Na^+^] of the pipette saline. Taken together, our FLIM recordings demonstrate that baseline [Na^+^]_i_ is rather uniform throughout the proximal apical dendritic tree of CA1 pyramidal neurons, averaging around 10 mM. Furthermore, they indicate that spontaneous, action-potential related neuronal activity and/or currents through TTX-sensitive Na^+^ channels cause Na^+^ influx that increases the apparent “baseline” [Na^+^]_i_ of dendrites.

### Line-scan imaging of dendritic [Na^+^]_i_ transients

While FLIM enabled quantitative determination of dendritic [Na^+^]_i_, its relatively long photon collection periods and low signal-to-noise ratio preclude studying rapid diffusion of Na^+^. We therefore switched to intensity-based line scan imaging of Na^+^ (Rose et al., 1999). Cells were filled with SBFI and held in the voltage-clamp mode while perfusing slices with TTX. To evoke a local increase in dendritic [Na^+^]_i_, a glutamate-filled iontophoresis pipette was positioned close (1-10 µm) to a primary dendrite (Fig. 1F). Next, a scan line was positioned onto the dendrite (Fig. 1F) and line scanning was performed at frequencies of 233-747 Hz for a total recording length of 5-18.5 sec. 500 ms after starting the scan, glutamate was applied for 100 ms.

Glutamate induced an inward current which started at 25 ms ± 21 ms after starting the iontophoresis (median=16 ms), reached its peak amplitude (mean 1304 ± 745 pA, median=1132 pA) at 210 ± 65 ms (median=214 ms) and had declined to 10% of its maximal amplitude after another 467 ms ± 190 ms (median=460 ms; n=18, N=18) (Fig. 1 F,G). In addition, glutamate caused a transient increase in [Na^+^]_i_, followed by a monotonic recovery to the initial baseline (Fig. 1F). Iontophoresis in the absence of glutamate in the pipette did not evoke membrane currents, nor significant changes in cellular fluorescence (n=7, N=7; not illustrated).

For an unbiased identification of the region of maximum Na^+^ influx, a Python-based automated analysis was employed, generating 10.4 µm (50 pixels) long “Sections Of Interest” (SOIs) on the scanned line. The region with the largest relative change in fluorescence, termed SOI 0, was generally located in the part of the dendrite closest to the tip of the iontophoresis pipette as well as in its direct projected ejection stream (Fig. 1F). Based on a baseline dendritic [Na^+^]_i_ of 5.3 mM in the presence of TTX (see above), [Na^+^]_i_ increases in SOI 0 had an average peak amplitude of 25.0 ± 10.1 mM (median=22.2 mM; n=18, N=18). Aligning the fluorescence changes in SOI 0 with the somatic inward current showed that [Na^+^]_i_ continued to rise for about 430 ms after commencing the glutamate application (426 ± 154 ms, median=393 ms; n=18, N=18). At this time point, the current had declined to about 50% of its maximal amplitude and reached a second turning point in its 1^st^ deviation, indicating slow cessation of [Na^+^]_i_ influx (Fig. 1G). Thus, inward currents were still ongoing whilst [Na^+^]_i_ already started to decay back to baseline, indicating a period of concomitant influx and clearance of Na^+^ from the site of influx (Fig. 1G).

To address the relevance of ionotropic glutamate receptors, slices were perfused with saline containing receptor blockers. As shown in Fig. 1H, both somatic inward currents and dendritic [Na^+^]_i_ transients were strongly and reversibly dampened (currents reduced to 16 ± 7% (median=15%), [Na^+^]_i_ transients reduced to 9 ± 6% (median=10%)) by perfusing slices with the NMDA-R blocker APV (100 µM) and the AMPA-R blocker NBQX (20 µM) (n=6, N=6; Fig. 1H).

The contribution of NKA mediated [Na^+^]_i_ extrusion on the clearance of dendritic Na^+^ elevations was probed for by performing experiments at 32 °C. At both room and elevated temperature, recovery from glutamate-induced Na^+^ transients followed a monoexponential decay with decay time constants of about 1.4 s (22°C: mean=1.42 s ± 0.65 s, median=1.26; n=18, N=18; 32 °C: mean=1.39 s ± 0.83 s, median=1.14; n=15, N=11; p=0.704) (Fig. 1 I, J).

The data therefore suggests, that (temperature-dependent) active transport processes, such as NKA mediated extrusion had virtually no effect on the clearance of [Na^+^]_i_ elevations in dendrites. Finally, we assessed the dependence of the amplitude of the Na^+^ load on the recovery rate, plotting the decay time constants against the respective peak [Na^+^]_i_ amplitudes (results from both temperatures pooled). The data shows no correlation between the two parameters (Pearson correlation < 0.01; Spearman correlation= 0.17) (Fig. 1K), again indicating that [Na^+^]_i_ elevations are cleared by a diffusive process.

Taken together, these experiments show that intensity-based line scanning enables measurement of glutamate-induced Na^+^ influx in dendrites. Furthermore, our results confirm the expected dominating role of ionotropic glutamate receptors in the generation of glutamate-induced currents and intracellular Na^+^-signals in CA1 neurons (Rose and Konnerth, 2001; Miyazaki and Ross, 2017). Finally, the data suggests that active transport processes do not play a major role in the clearance of local Na^+^-increases, but instead point towards lateral diffusion as a major contributor.

### Analysis of the lateral spread of Na^+^ within primary dendrites

For unbiased analysis of the lateral spread of dendritic Na^+^ signals in line scan recordings, we employed our Python-based tool including automated definition of SOIs at a standard length 5.1 µm/25 pixels, as well as automated detection of peak amplitudes in these SOIs. Fig. 2A-C illustrates a typical experiment performed in a primary dendrite (n=18 cells, N=18 slices). As described above, SOI 0 generally corresponded to the segment closest to the application pipette. For further analysis, we normalized peak amplitudes and peak time points of neighboring SOIs to that of the respective SOI 0.

**Figure 2:**
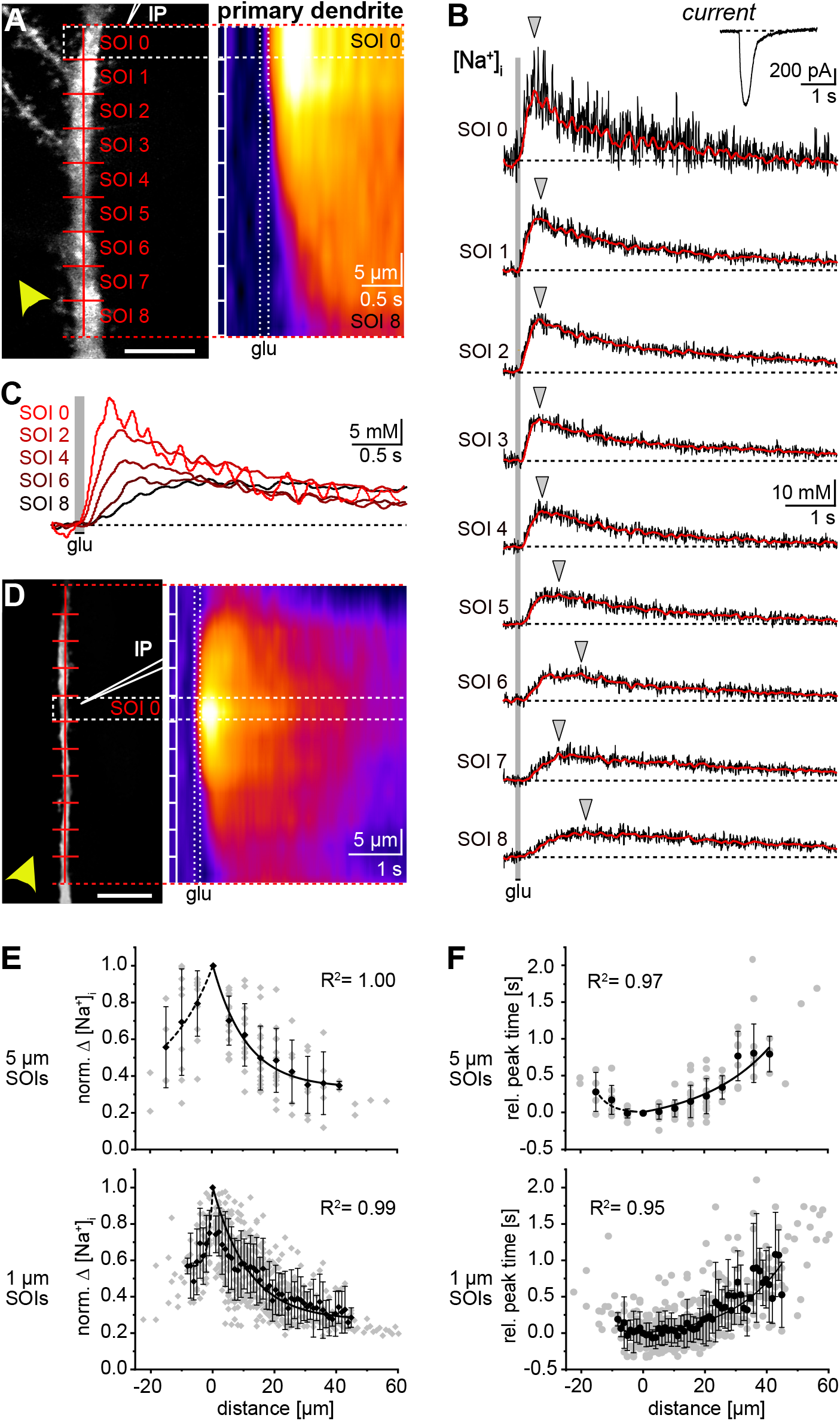
Spread of Na^+^ along primary dendrites. A: Projection of an SBFI-filled primary dendrite. Red line: position of the scan line; with SOIs spanning 5.1 µm each. IP: iontophoresis pipette; arrowhead indicates flow of bath perfusion. Scale bar: 10 µm. Right: False color-coded (x,t) image of the baseline-corrected line scan with SOI 0 outlined. Vertical boxes indicate SOIs, dotted box indicates glutamate application (100 ms). B: Glutamate-induced [Na^+^]_i_ transients in SOIs 0-8. Black: baseline corrected traces (subjected to 50 Hz low pass FFT), red: filtered traces as given by the Python-based program. Grey area indicates glutamate application. Grey triangles indicate the peak time points. Trace on the top right shows the somatic current. C: Overlaid traces of Na^+^ transients from SOI 0, 2, 4, 6 and 8 at higher time resolution. D: Projection of an SBFI-filled primary dendrite subjected to imaging where SOI 0 is outlined. IP: iontophoresis pipette; arrowhead indicates flow of bath perfusion. Scale bar: 10 µm. Right: False color-coded (x,t) image of the baseline-corrected line scan with SOI 0 outlined; dotted rectangular box indicates glutamate application (100 ms). E: Data of 18 experiments showing normalized peak changes in [Na^+^]_i_ (norm. Ω[Na^+^]_i_) versus distance from SOI 0 for a SOI length of 5.1 µm (Top) and of 1.02 µm (Bottom). Negative distances indicate dendritic sections within the perfusion direction relative to SOI 0. Shown are individual data points (grey symbols), means (black symbols) and standard deviations (whiskers). Black dotted line shows exponential fit for SOIs within perfusion direction (R^2^=0.99), black line for SOIs against the perfusion flow (R^2^=1.00). F: Peak time points after stimulation versus distance from SOI 0; same data set and illustration as shown in E.

We found that peak amplitudes of [Na^+^]_i_ transients declined with increasing distance from SOI 0 along the dendrite, while the latency from the stimulation onset to the peak time point increased (Fig. 2 A-D). At a distance of 40 µm from the stimulation site, [Na^+^]_i_ transients had declined to 30-40% of the initial peak amplitude and their peak was reached after about 1 sec (Fig. 2 E,F). In a subset of recordings (n=4, N=4), dendrites could be recorded for a length of at least 15 µm both up- and downstream of the pipette tip (Fig. 2D). Here, the instant decay in peak amplitude, together with the increase in the latency, was detected for both directions, i. e., in SOIs along and against the main perfusion direction (Fig. 2 D-F). This observation suggests that dendritic [Na^+^]_i_ transients detected upstream and downstream of SOI 0 were not primarily caused by perfusion-driven (asymmetrical) diffusion of glutamate in the extracellular space, but predominately due to intracellular spread of [Na^+^]_i_ from its initial site of influx. Moreover, reducing the length of SOIs to 1.02 µm (5 pixels) showed that the decay in peak amplitude was already apparent in the second segment (i.e. after 1.02 µm), indicating that maximum Na^+^ influx was confined to a small section of the dendrite (Fig. 2 E,F).

In summary, these results demonstrate that upon local Na^+^ influx caused by activation of ionotropic glutamate receptors, [Na^+^]_i_ rapidly spreads up- and downstream along primary apical dendrites. [Na^+^]_i_ transients are still detectable at distances of 40-50 µm, at which their peaks are delayed by about 1 sec.

### Morphological characteristics of primary and secondary dendrites

Dendritic spines can transiently trap a certain fraction of diffusing ions and molecules as shown for Cl^-^ and IP_3_ (Santamaria et al., 2006, 2011; Mohapatra et al., 2016). To evaluate the dependence of the spread of Na^+^ on the morphological properties and spine density of dendrites, we analyzed 3D-reconstructed images of SBFI-filled apical dendrites at three different times in culture (DIV 3-6, 10-25, 35-50). At DIV 10-25, primary dendrites at a distance of 10-60 µm from the soma had an average diameter of 1.8 ± 0.2 µm (median=1.7 µm; n=18, N=18) (Fig. 3A, B). Secondary dendrites emerged from these primary dendrites at a distance of 40-60 µm from the soma. After DIV 10-25, these secondary dendrites had a significantly smaller average diameter of 1.3 ± 0.1 µm (median=1.3 µm; n=18, N=18; p=5.45E-10). The same was true for secondary dendrites at DIV 3-6 (mean=1.2 ± 0.1 µm, median=1.2 µm; n=17, N=17; p=5.40E-10) and at DIV 35-50 (mean=1.2 ± 0.2 µm, median=1.2 µm; n=13, N=13; p=1.58E-8) (Fig. 3A, B). Taken together, this data shows that secondary dendrites after different times in organotypic culture exhibited a similar diameter. Moreover, the diameter of secondary dendrites was significantly smaller than the diameter of primary dendrites.

**Figure 3:**
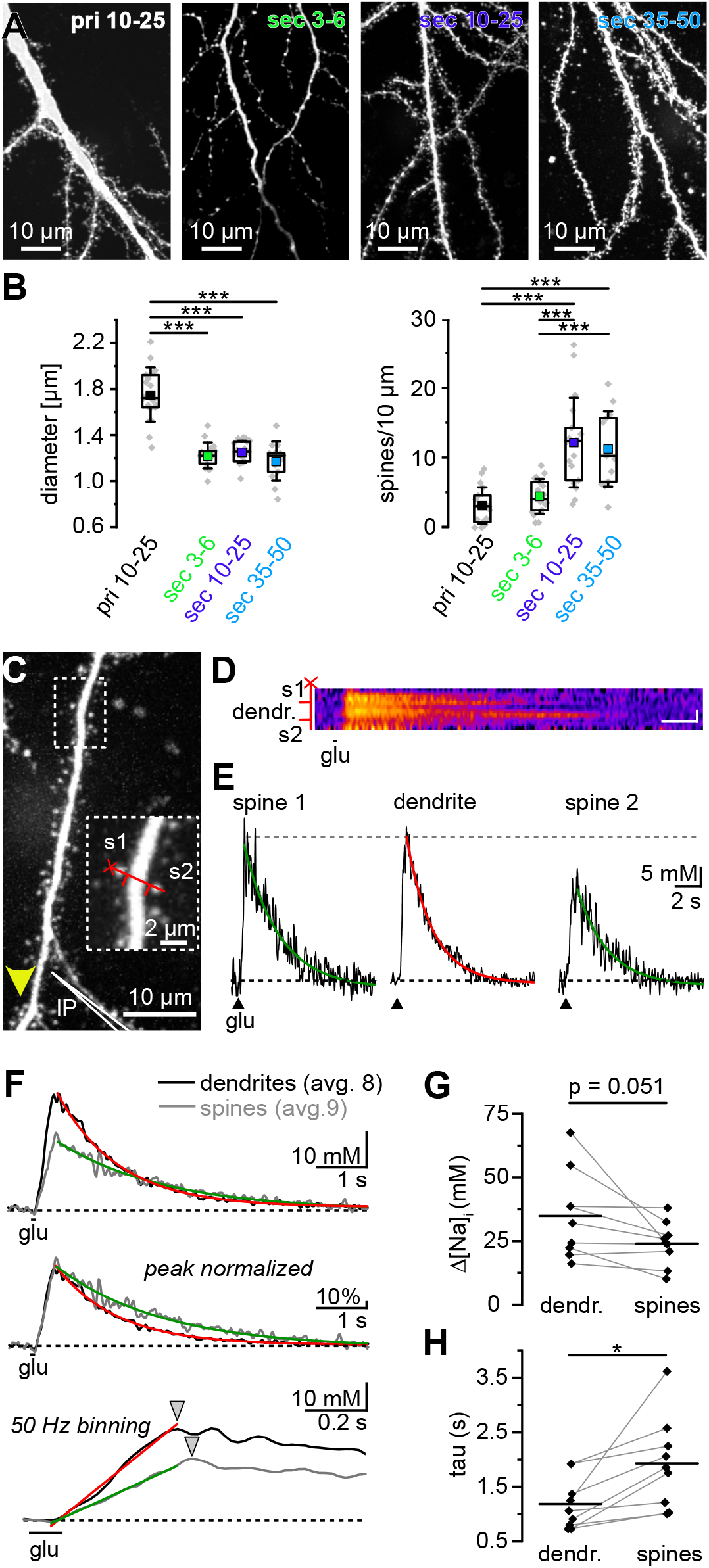
Morphological characteristics of dendrites and diffusion of Na^+^ from dendrites into spines. A: Maximal projections of primary (pri) and secondary (sec) dendrites filled with SBFI in slices cultured for 3-6, 10-25 or 35-50 days. B, C: Boxplots showing dendrite diameter (left) and spine density (right) of primary dendrites (DIV 10-25; n=18, N=18) and of secondary dendrites at DIV 3-6 (n=17, N=17), DIV 10-25 (n=18, N=18) and DIV 35-50 (n=13, N=13). Shown are individual data points (diamonds), means (squares), medians (horizontal lines), quantiles (boxes), and standard deviations (whiskers). For statistical analysis, normal distribution of the data was determined by the Shapiro Wilk test and one-way ANOVA with Bonferroni *post hoc* analysis was used for normal distributed data. Statistical significance is indicated by asterisks: ***p<0.001. C: Maximal projection of an SBFI filled spiny dendrite at DIV 10-25. IP: iontophoresis pipette; arrowhead indicates flow of bath perfusion. Inset: Section shown at higher magnification, indicating the placement of the line scan (red line) over the dendrite and two adjacent spines (s1, s2). D: False color image of a line scan, taken from the dendrite shown in A with a binning of 2 pixels. Black line below indicates the glutamate application (100 ms). Scale bars represent the following: Horizontal scale bar 1 s, vertical scale bar 1 µm. E: Na^+^ transients taken from the line scan shown in D, filtered as given by the Python-based program. The recovery to baseline was fitted monoexponentially (colored lines). F: Top. Averaged traces taken from dendrites (n=8, N=8; black trace) and spines (n=9, N=8; grey trace), after binning individual measurements to 50 Hz. Colored traces represent monoexponential fits of the decay. Center: Peak-normalized traces. Bottom: traces at higher temporal resolution. Linear fits depict the slopes of the rising phase. Arrowheads point at the peak time points. G, H: Histograms showing peak changes in [Na^+^]_i_ (Ω[Na^+^]_i_) and decay time constants (τ) of the recovery from Na^+^ transients within dendrites (n=8, N=8) and directly adjacent spines (n=9, N=8). Shown are individual data points (black diamonds) and means (short horizontal lines). Data points derived from a given dendrite and adjacent spine are connected by grey lines. Normal distribution of the data was determined by the Shapiro Wilk test; paired sample t-tests were used to determine the statistical significance between data sets and is indicated as: 0.05<*p<0.01.

Primary dendrites at DIV 10-25 exhibited an average spine density of 3 ± 3 spines per 10 µm length (median=3 spines; n=18, N=18). Spine density was significantly higher in secondary dendrites at DIV 10-25, which displayed 12 ± 6 spines/10 µm (median=12 spines; n=18, N=18; p=3.49E-6), and which was similar to DIV 35-50 secondary dendrites (11 ± 5 spines, median=10 spines; n=13, N=13; p=0.68). Secondary dendrites at DIV 3-6, on the other hand, showed a density of 4 ± 2 spines (median=4 spines; n=17, N=17), which was comparable to spine densities in DIV 10-25 primary dendrites (p=0.13), but significantly lower than in DIV 10-25 (p=5.55E-5) and DIV 35-50 secondary dendrites (p=8.45E-5) (Fig. 3A, B).These results show that while spine densities were at a low level and comparable between primary dendrites at DIV 10-25 and secondary dendrites at DIV 3-6, they were significantly higher in secondary dendrites at longer culturing periods (DIV 10-25 and DIV 35-50).

### Spread of Na^+^ signals from dendrites and into dendritic spines

In a next step, we analyzed if Na^+^ signals also spread from dendrites into adjacent spines. Experiments were done in slices at DIV 10-25, i.e., at a time point when spine densities reached a high and stable level. To this end, we positioned a line scan perpendicular to a dendrite to cover a spine head visually separated from it (Fig. 3C). To exclude an eventual direct stimulation of glutamate receptors on spines, experiments were performed at distances of >10 µm upstream of the iontophoresis pipette (Fig. 3C). SOIs were then defined manually before performing Python-based automated analysis for baseline correction and filtering of the resulting transients. Further analysis, which included the determinations of peaks and decay time constants (τ) were again commenced manually using the Origin software.

We found that Na^+^ transients were not only observed in distant dendritic sections, but also in spine heads at these sites (n=8 dendrites/9 spines, N=8) (Fig. 3 D-E). Interestingly, peak amplitudes of spine head Na^+^ transients were either comparable (4/9 spines) or lower (5/9 spines) than those of their directly adjacent parent dendrites (Fig. 3 D-E). On average, amplitudes within spine heads amounted to 24.0 ± 8.7 mM (median=24.5 mM; n=9, N=9) compared to 34.5 ± 18.3 mM in dendrites (median=28.2 mM; n =8, N= 8; p=0.051) (Fig. 3G). Binning scans to 50 Hz and averaging traces for all dendrites and all spines revealed that dendritic Na^+^ signals had a steeper rise and that peak amplitudes were reached about 40 ms earlier in dendrites than in adjacent spine heads (Fig. 3F). While both dendrites and spines followed a monoexponential recovery to baseline, decay time constants were significantly larger in spines (1.93 ± 0.84 s, median=1.85 s) as compared to those in directly adjacent parent dendrites (1.09 ± 0.41 s, median=1.0 s; p=0.012) (Fig. 3 F,H).

These results show that Na^+^ readily spreads from dendrites into adjacent spines. At the same time, the lower average amplitudes and slower kinetics of spine Na^+^ signals indicate that spine necks represent an anatomical and/or functional diffusion barrier for Na^+^.

### Spread of Na^+^ signals within secondary dendrites at different DIV

Having established that diameter and spine density differ between primary and secondary dendrites at different DIV, we next analyzed the lateral spread of Na^+^ signals along secondary dendrites, using the same strategy as described above. As expected, x-t-images revealed a transient increase in dendritic [Na^+^]_i_ along secondary DIV 10-25 dendrites in response to glutamate (n=18, N=18) (Fig. 4A). Amplitudes in DIV 10-25 secondary dendrites were higher than those observed in primary dendrites with an average Ω[Na^+^]_i_ 41.5 ± 18.5 mM (median: 36.9 mM; n=18; N=18; p=0.004). Similar to what was observed in primary dendrites, [Na^+^]_i_ declined monoexponentially with increasing distance from SOI 0, while the latency of [Na^+^]_i_ changes followed a monoexponential increase (Fig. 4 A-D). The same behavior was observed in secondary dendrites at DIV 3-6 (n=17, N=17) (Fig. 4E) and DIV 35-50 (n=13, N=13) (Fig. 4F).

**Figure 4:**
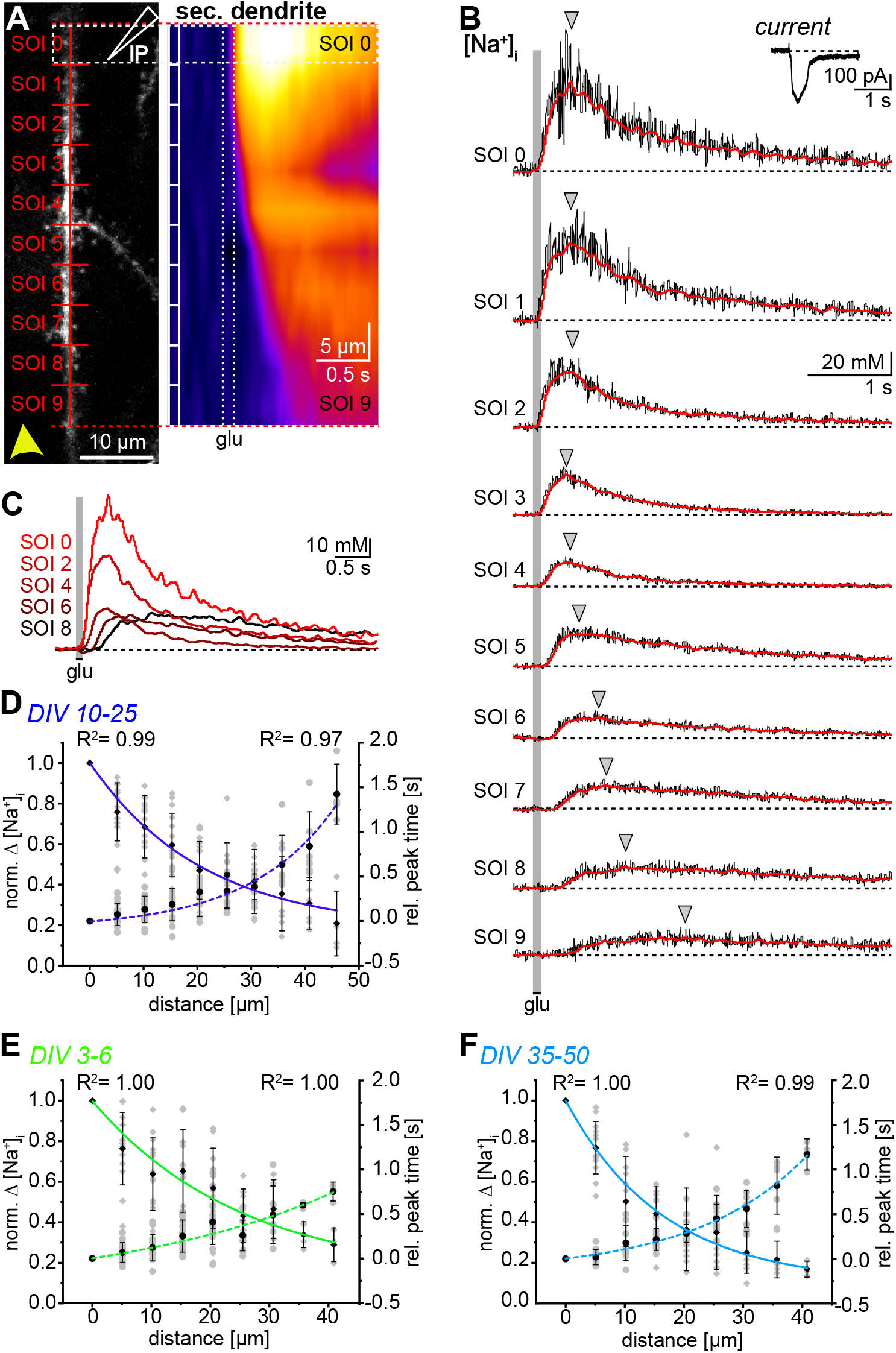
Spread of Na^+^ along secondary dendrites. A: Projection of an SBFI-filled secondary dendrite. The red line indicates the scan line, SOIs are 5.1 µm each. IP: iontophoresis pipette; arrowhead indicates flow of bath perfusion. Right: False color-coded (x,t) image of the baseline-corrected line scan. Vertical boxes indicate SOIs with SOI 0 outlined, the dotted rectangular box indicates the glutamate application (100 ms). B: Glutamate-induced [Na^+^]_i_ transients in SOIs 0-9. Somatic current is shown on the top right. Black: baseline corrected traces (subjected to 50 Hz low pass FFT), red: filtered trace as given by the Python-based program. Grey area indicates glutamate application. Grey triangles indicate the peak time points. C: Overlaid traces from SOI 0, 2, 4, 6 and 8 at higher time resolution. D, E, F: Normalized peak changes in [Na^+^]_i_ (norm. Ω[Na^+^]_i_; diamonds) and peak time points after stimulation (circles) versus distance from SOI 0 in DIV 10-25 secondary dendrites (n=18, N=18), DIV 3-6 secondary dendrites (n=17, N=17) and DIV 35-50 secondary dendrites (n=13, N=13). Shown are individual data points (grey symbols), means (black symbols) and standard deviations (whiskers). Colored lines represent monoexponential fits of the data, with corresponding R^2^ values indicated (full line: Ω[Na^+^]_i_; dotted line: peak time points).

To compare the decay in amplitude and the increase in the latency of Na^+^ signals between different dendrite order and DIV, we tested the correlation between the before shown monoexponential fits taken from primary dendrites at DIV 10-25 (Fig. 2 E,F) and secondary dendrites at DIV 3-6, DIV 10-25 and DIV 35-50 (Fig. 4 D-F). Calculation of both the Pearson and Spearman correlations revealed that monoexponential fits for all secondary dendrites showed a correlation >0.99. Furthermore, a strong correlation was found between secondary and primary dendrites (DIV 10-25; correlation coefficients >0.97). This indicates that the lateral spread of Na^+^ was similar in all preparations investigated.

In summary, these data demonstrate that local [Na^+^]_i_ transients efficiently spread along secondary dendrites, exhibiting a monoexponential decay in peak amplitudes and a monoexponential increase in latency with distance from the stimulation site, similar to what was observed in primary dendrites. Moreover, our results indicate that the spread of Na^+^ is largely independent from the time in culture and thus from the diameter of dendrites. Finally, we conclude that Na^+^ diffusion is independent from spine density (which tripled on secondary dendrites from DIV 3-6 to DIV 10-50).

### Determination of apparent diffusion coefficients of Na^+^ in dendrites

For determination of the apparent diffusion coefficient (*D*_*app*_) of Na^+^ along dendrites over time, we employed an analysis similar to that first proposed by Santamaria et al. (Santamaria et al., 2006). For this, we utilized the Python-based program to establish 1.02 µm SOIs (5 pixels) along the line-scanned dendrites and performed a temporal binning to 50 Hz. All profiles had their peak aligned to SOI 0 which was defined as the origin, *x*_*o*_*=0* (Fig. 5A). For further analysis, we normalized the data using Eq. 1 (see Materials and Methods) and pooled the data taken from either condition (DIV 10-25 primary dendrites and DIV 3-6, DIV 10-25, DIV 35-50 secondary dendrites) (Fig. 5B). We then calculated the change in variance ⟨*x*^*2*^_*exp*_⟩ over time using Eq. 3 (Fig. 5B) and finally *D*_*app*_ using Eq. 4 (Fig. 5C) for all four experimental conditions.

**Figure 5:**
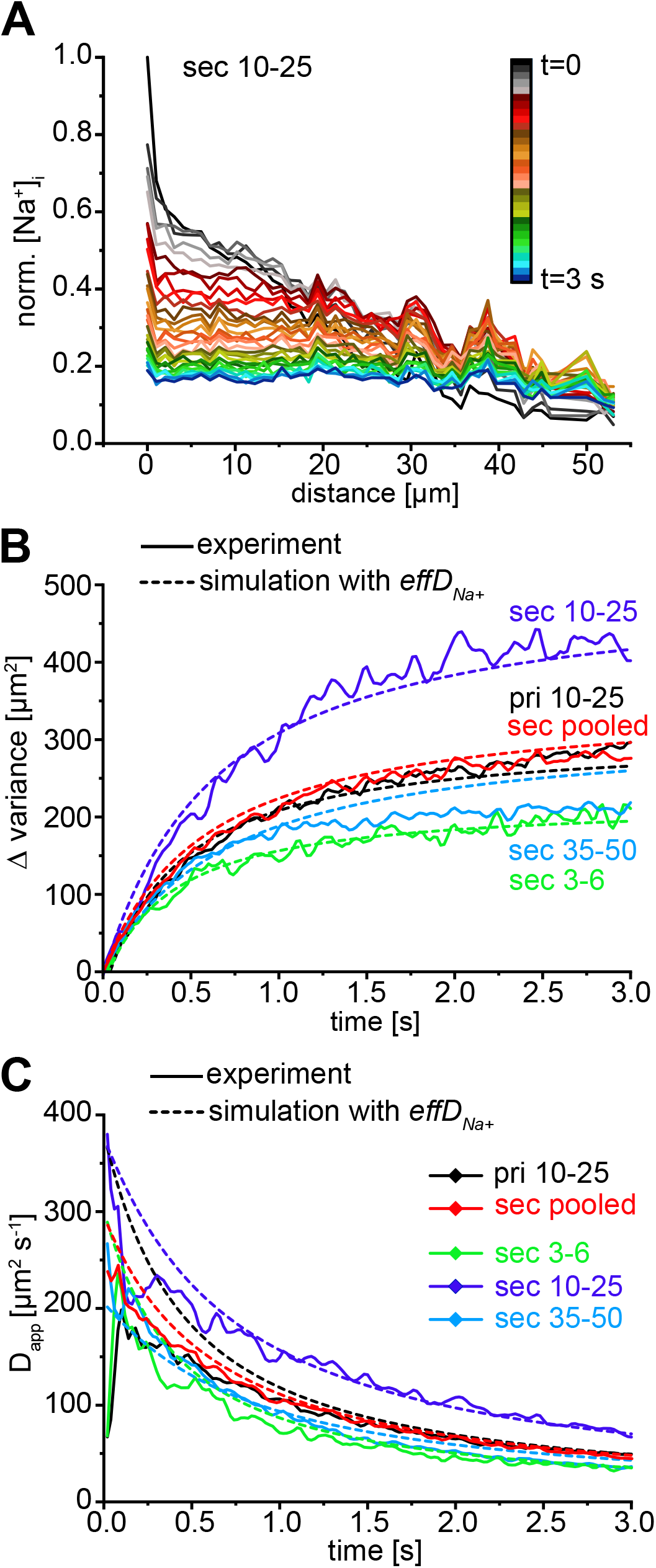
Impact of dendrite morphology on the diffusional spread of Na^+^. A: Spatial profiles of normalized changes in [Na^+^]_i_ for all investigated DIV 10-25 secondary dendrites. Shown are concentration profiles taken every 100 ms (10 Hz). Color code depicts the point in time at which spatial [Na^+^]_i_ distributions were determined. B: Plots showing changes in variance over time in DIV 10-25 primary dendrites (black), in secondary dendrites at DIV 3-6 (green), DIV 10-25 (dark blue) and DIV 35-50 (light blue), and for all secondary dendrites (pooled data, red), calculated from profiles such as shown in A. Dotted lines illustrate the mathematical analysis of the diffusional spread using the boundary effects and initial conditions provided by the respective conditions employing the for each condition. C: 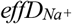 Apparent Na^+^ diffusion coefficients (D_app_) resulting from variances depicted in B. Dotted lines illustrate the analysis using the boundary effects and initial conditions employing the 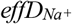 for each condition.

The analysis of ⟨*x*^*2*^_*exp*_⟩ suggested similar calculated *D*_*app*_ for all conditions exhibiting initial values of 200-400 µm^2^/s, which declined to about 50-100 µm^2^/s within 2-3 seconds, indicating a slowing down of diffusional dynamics (Fig. 5C). Moreover, the analysis showed that the initial distribution of the [Na^+^]_i_ had a large variance ⟨*x*^*2*^_*exp*_⟩_*o*_ in the range of 200-400 µm^2^ (Table 1). Of note, *D*_*app*_ is a measure valid only for the specific experimental conditions. As a next step, we therefore included boundary conditions in our calculation, more specifically the length of the dendritic segment monitored and the width of the initial [Na^+^]_i_ distribution determined at *t=0*. To further evaluate the experimentally determined trajectory of *D*_*app*_ over time, we thus quantified the effects of the finite length of the line scan and the initial spread of the Na^+^ signal on the estimation of *D*_*app*_, assuming that Na^+^ diffusion was unhindered.

To do so, we simulated a normal 2D diffusion process for each condition using Eq. 5-7. We thereby shifted *t*_*D*_ in Eq. 5 to account for the initial condition, *t*_*D*_ = *t* + *t*_*o*_, with *t* being the line scan time. For each simulation, we used the corresponding line scan length and sampling settings and determined the most effective 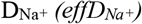 (Table 1). All conditions showed that the simulation of an unhindered, normal diffusive process led to a trajectory of ⟨*x*^*2*^_*exp*_⟩ which resembled those obtained experimentally (Fig. 5B, dotted lines). This shows that the dynamics measured experimentally follow characteristics of normal diffusion. Moreover, we found that using the Na^+^ diffusion coefficient, which was determined earlier in muscle cells 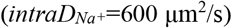 (Kushmerick and Podolsky, 1969), also fit all conditions with an error of <5%, indicating that Na^+^ diffusion is not significantly different between the different dendrites investigated. It is noteworthy that this is the case for all conditions, although [Na^+^]_i_ loads were significantly higher in secondary as compared to primary dendrites. Thus, the data again shows that the diffusive dynamics and therefore also the diffusion coefficient is not dependent on the amplitude of the Na^+^ load.

Finally, the resulting profiles were analyzed with the same procedure we used to determine *D*_*app*_ (Eq. 2-4; Fig. 5C, dotted lines). The resulting simulated plots corresponded to the experimentally derived data, also showing a decline in *D*_*app*_ from initially 200-400 µm^2^/s to about 50-100 µm^2^/s within 2-3 seconds.

Taken together, the data demonstrates that the lateral Na^+^ diffusion over time was similar between the different dendrites and conditions analyzed. Our results also show that Na^+^ diffusion was a) independent of dendrite diameter and b) independent of spine density in this preparation. Moreover, we found that the lateral spread of Na^+^ in dendrites follows normal diffusion and is well described using a diffusion coefficient of 600 µm^2^/s.

### Modeling of Na^+^ dynamics in dendrites

To further understand intra-dendritic Na^+^ dynamics, we simulated Na^+^ movement throughout a modeled dendrite of a pyramidal neuron (Fig. 6A). We increased [Na^+^]_i_ in a ~5 µm segment of the dendrite (arrows in Fig. 6 A, B) for 500 ms, and used 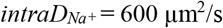 to examine the lateral spread of Na^+^ through the dendrite (Fig. 6B). Line scan images showed a decrease and delay of the signal with increasing distance from the stimulated segment with characteristics similar to those observed experimentally (Fig. 6C). Concentration profiles taken at the peak of the signal, as well as every 20 ms over the course of 3 s, showed a reduction of the amplitude, as well as a spreading of Na^+^ over time (Fig. 6D), again reflecting the kinetics observed in experiments. Finally, Na^+^ transients recorded at 0, 5.8, 9.2, 12.1, 15.5 and 22.2 µm from the stimulation site showed trajectories similar to those observed in experiments (Fig. 6E).

**Figure 6:**
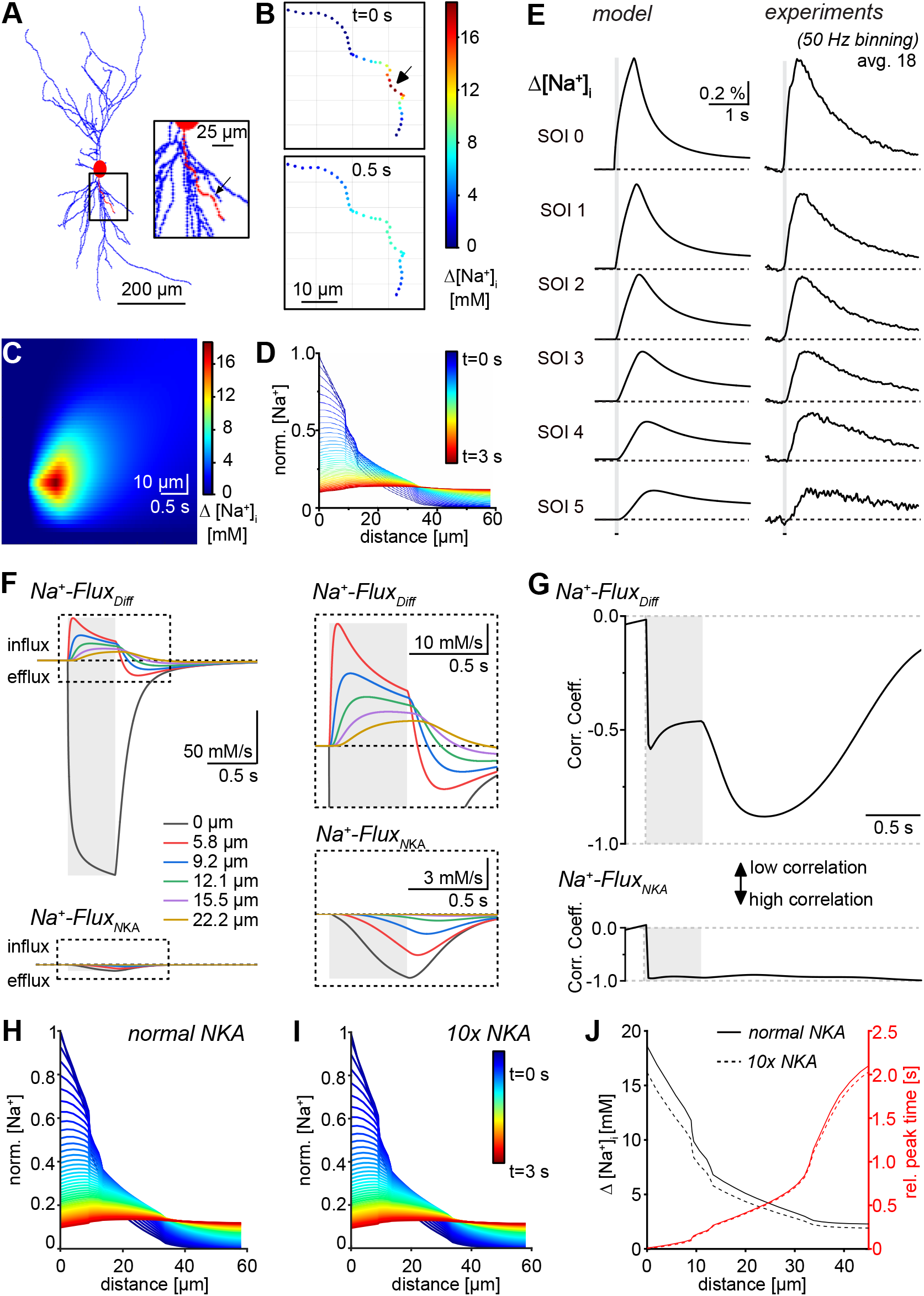
Modeling of Na^+^ dynamics in dendrites. A: 2-Dimensional projection of the measured CA1 pyramidal neuron (NeuroMorpho.org; (Buchin et al., 2022) NMO_276156; (Tecuatl et al., 2024). The soma is indicated as large red oval, the dendrite used for the simulation is highlighted in red (also see Insert). The arrow points to the dendrite segment subjected to an increase in Na^+^. B: Color-coded images, showing the [Na^+^]_i_ in the dendrite at t=0 s and t=0.5 s after reaching the maximum peak at the segment associated with the highest Na^+^ amplitude (at 0 µm; indicated by an arrow). C: Image of the color-coded line scan, showing the [Na^+^]_i_ spread through the dendrite over time. D: Spatial concentration profiles of normalized changes in [Na^+^]_i_ taken every 20 ms. Color-code depicts the point in time at which [Na^+^]_i_ distributions were determined. E: Left: [Na^+^]_i_ transients in the stimulated segment (0 µm), as well as segments at 5.8, 9.2, 12.1, 15.5 and 22.2 µm distance. Right: Corresponding experimental data from primary dendrites (n=18, N=18; see Fig. 2). Individual traces were averaged and normalized to the peak at 0 µm distance. The grey column indicates the duration of the glutamate application. F: Na^+^ fluxes at the before mentioned distances from the stimulated site, including diffusion fluxes (*Na*^*+*^*-Flux*_*Diff*_) and NKA-mediated extrusion fluxes (*Na*^*+*^*-Flux*_*NKA*_). Inserted boxes show blown up images of the traces on the left. G: Correlation coefficient of [Na^+^]_i_ transients with *Na*^*+*^*-Flux*_*Diff*_ and *Na*^*+*^*-Flux*_*NKA*_ over time. Note that correlation coefficient of 0 and −1, respectively, indicate no correlation and strong correlation between [Na^+^]_i_ and respective flux. H, I: [Na^+^]_i_ normalized with respect to the peak value at the stimulated segment as a function of distance from the stimulated segment at different times (color-coded) using normal NKA-mediated clearance (H) and enhanced (10 times that of normal) NKA-mediated clearance (I). J: Peak [Na^+^]_i_ (black) and time of the peak (red) as functions of distance from the stimulated segment at normal NKA-mediated clearance (solid lines) and enhanced (10 times that of normal) NKA-mediated clearance (dashed lines).

Next, we addressed the relevance of Na^+^ diffusion flux (*Na*^*+*^*-Flux*_*Diff*_) versus NKA-mediated extrusion (*Na*^*+*^*-Flux*_*NKA*_) (Fig. 6F). Note that this analysis did not include the influx of Na^+^ into the dendrite, but the dynamics after entry. The data shows that upon stimulation, Na^+^ quickly diffuses out of the segment subjected to a [Na^+^]_i_ increase (0 µm) into neighboring segments (Fig. 6F, top panels). Following segments typically show a net influx of Na^+^ followed by a net efflux, whereby the degree of both decreases with increasing distance from the stimulation site (Fig. 6F). This indicates the diffusional process and the diminishing of Na^+^ gradients at segments which show small Na^+^ amplitudes. In contrast, NKA makes only a small contribution to Na^+^ efflux in all segments (Fig. 6F, bottom panels). Moreover, the model demonstrates that while diffusion occurs rapidly and directly following the onset of Na^+^ entry, *Na*^*+*^*-Flux*_*NKA*_ exhibits significantly slower kinetics as it increased steadily during the rising phase of the Na^+^ signal. Therefore, the traces also show that NKA-mediated extrusion is governed by the [Na^+^]_i_ as it increases during transient [Na^+^]_i_ elevation.

To further investigate the relative influence of *Na*^*+*^*-Flux*_*Diff*_ and *Na*^*+*^*-Flux*_*NKA*_ on the dynamics of [Na^+^]_i_, we determined correlation coefficients between the modeled Na^+^ transients and these fluxes. The model shows a strong negative correlation between Na^+^ transients and *Na*^*+*^*-Flux*_*Diff*_ after stimulation, indicating a strong diffusional process (Fig. 6G, top panel). During the rising phase, the correlation between the [Na^+^]_i_ and Na^+^ diffusion increases only slightly, as the diffusion is only relevant in few segments in the early phase of the signal. After the maximum [Na^+^]_i_ is reached, the correlation increases further as diffusion occurs in all segments as Na^+^ spreads throughout the entire dendrite. With [Na^+^]_i_ approaching baseline, the correlation returns back to 0, indicating that Na^+^ diffusion decreases throughout the dendrite. This coincides with Na^+^ transients reaching similar values in all segments, leading to smaller Na^+^ gradients. The model also shows a high correlation between the Na^+^ transients and NKA mediated extrusion fluxes at all times, again indicating that NKA activity follows the [Na^+^]_i_ (Fig. 6G, bottom panel). It is noteworthy, that the NKA mediated fluxes are comparably smaller than the diffusional fluxes and thereby have a smaller effect on the fast clearance of Na^+^ throughout the dendrite.

In summary, using 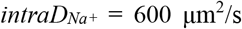 in the model results in kinetics of Na^+^ transients similar to those observed experimentally. These simulations also underline that lateral diffusion is key for the clearance of local [Na^+^]_i_ increases in early stages of the signal. Moreover, the data shows that once Na^+^ spreads throughout the dendrite and dendritic [Na^+^]_i_ equilibrates, diffusional gradients diminish and net diffusion is severely decreased. The extrusion of the Na^+^ must therefore be conducted through further transport processes, namely the NKA. We remark that while increasing the density of NKA (*ρ*) decreases the peak [Na^+^]_i_ at a given segment, the diffusion remains largely unaffected (Fig. 6 H-I). To see a noticeable effect on the speed at which the [Na^+^]_i_ peak propagates along the dendrite, one would have to increase the NKA density by at least an order of magnitude. Even increasing NKA density by a factor 10, the time (from the stimulation time) of the peak [Na^+^]_i_ remains unchanged as far as 35 µm from the stimulated segment (Fig. 6J). At larger distance from the stimulated segment, the time of the peak decreases slightly (e.g., 2.03 sec at normal NKA density vs 1.96 sec at 10 times higher density at 43.34 µm from the stimulating segment) (results not shown).

## Discussion

### Spread of Na^+^ within spiny dendrites

Na^+^-FLIM (Meyer et al., 2022) revealed that baseline dendritic [Na^+^]_i_ is ~10 mM and therefore in the range of [Na^+^]_i_ reported for somata of CA1 neurons (Diarra et al., 2001; Langer and Rose, 2009; Kelly and Rose, 2010; Azarias et al., 2013; Mondragao et al., 2016; Meyer et al., 2022). Moreover, we found that [Na^+^]_i_ was uniform in primary and secondary dendrites. This is similar to dendritic Ca^2+^ concentrations of CA1 neurons (Zheng et al., 2015), while in cortical pyramidal neurons, dendritic Cl^-^ concentrations slowly decline with increasing distance from the soma (Weilinger et al., 2022).

Local increases in dendritic [Na^+^]_i_ were induced by glutamate application for 100 ms, resulting in Na^+^ influx through ionotropic glutamate receptors (Rose and Konnerth, 2001; Mondragao et al., 2016; Miyazaki and Ross, 2017). This paradigm generated [Na^+^]_i_ signals large enough to be tracked over a longer distance along the dendrite. The mean amplitude of [Na^+^]_i_ signals in primary dendrites was 25 mM, but some [Na^+^]_i_ transients were also much larger. Earlier work has shown that brief suprathreshold (4-5 APs), local afferent stimulation caused a [Na^+^] increase of ~13 mM in CA1 dendrites, and ~35 mM in presumed active spines (Rose & Konnerth, J Neurosci 2001). Moreover, an LTP induction protocol (100 Hz/1 s) induced [Na^+^]_i_ increases by as much as 40-50 mM (Rose and Konnerth, 2001). Notably, Miyazaki and Ross (Miyazaki and Ross, 2017) demonstrated that with one subthreshold EPSP only, there is an increase in spine [Na^+^]_i_ by 5-6 mM, and in the adjacent dendrite by ~3 mM.

These results show that neurons experience significant [Na^+^]_i_ loads, particularly following temporal summation of EPSPs or during periods of intense simultaneous activity.

Peak amplitudes decayed and latencies increased monoexponentially with increasing distance to the influx site, indicating intracellular spread of Na^+^ due to diffusion towards unstimulated regions. This is in line with earlier studies performed in dendrites (Rose and Konnerth, 2001; Mondragao et al., 2016; Miyazaki and Ross, 2017), axons (Kole et al., 2008; Fleidervish et al., 2010; Baranauskas et al., 2013; Filipis and Canepari, 2021), or astrocytes (Langer et al., 2012; Langer et al., 2017; Moshrefi-Ravasdjani et al., 2017).

Our line-scan measurements showed that dendritic Na^+^ signals invade adjacent spines, confirming results obtained upon induction of back-propagating action potentials (Rose et al., 1999). Fast diffusive movement of Na^+^ also occurs from spines into dendrites (Rose and Konnerth, 2001; Miyazaki and Ross, 2017, 2022). Here, spine Na^+^ signals generated by influx from dendrites were either similar or smaller in amplitude than those of parent dendrites. Moreover, their rise and decay times were longer, suggesting that spine necks represent a diffusional barrier for Na^+^, as reported for other molecular species, including Ca^2+^ (Yuste and Denk, 1995; Sabatini et al., 2002). Spines can also undergo independent Ca^2+^ signaling from parent dendrites which mainly results from the presence of Ca^2+^ buffers (Sabatini et al., 2002). Molecular diffusion between spines into dendrites is restricted by spine necks, for which electro-diffusion models need to be employed owing to their small diameter (Qian and Sejnowski, 1989; Holcman and Yuste, 2015; Cartailler and Holcman, 2018).

### Determination of Na^+^ diffusion coefficients in dendrites

A main goal was to determine dendritic D_Na+_ and to reveal the influence of spines on Na^+^ diffusion. For aqueous solutions, a D_Na+_ of 1,200-1,500 µm^2^/s was reported (Kushmerick and Podolsky, 1969; Pusch and Neher, 1988; Lobo, 1993). Values for intracellular diffusion reach from 600 µm^2^/s for muscle cells (Kushmerick and Podolsky, 1969), to 790 µm^2^/s for oocytes (Allbritton et al., 1992), to 1300 µm^2^/s along lizard axons (David et al., 1997). To determine dendritic D_Na+_, we followed an approach employed earlier (Santamaria et al., 2006, 2011; Mohapatra et al., 2016), combining imaging with computer simulations.

High-affinity Ca^2+^ indicators like Fura-2 or Oregon-Green mediate considerable buffering and thereby distort Ca^2+^ signals (Neher and Augustine, 1992; Zhou and Neher, 1993; Maravall et al., 2000). Notably, this is not the case for Na^+^ signals reported by Na^+^ indicators. SBFI displays a K_d_ of ~24 mM (Donoso et al., 1992; Jung et al., 1992; Rose et al., 1999; Sheldon et al., 2004; Meier et al., 2006), and does not buffer Na^+^ in the concentration range employed (Sabatini et al., 2001; Mondragao et al., 2016; Canepari and Ross, 2024). Furthermore, no relevant endogenous Na^+^ buffer systems exist (Despa and Bers, 2003; Fleidervish et al., 2010; Canepari and Ross, 2024). The decrease in dendritic D_Na+_ as compared to that in aqueous solutions is probably mostly due to intracellular organelles. Finally, as the diffusional mobility of SBFI will presumably not exceed 10-15 µm^2^/s (Yuste et al., 2000), the D_Na+_ determined here (600 µm^2^/s) will most likely reflect mobility of Na^+^, and not of SBFI.

Analysis of the spatial variance of dendritic [Na^+^]_i_ changes over time showed a non-linear relationship in all preparations. Similarly, *D*_*app*_ decreased non-linearly, indicating a slowing down of Na^+^ diffusion with time and distance. Earlier work has attributed these phenomena to a trapping of molecules and ions by spines, resulting in anomalous diffusion during first 1000 ms after influx (Santamaria et al., 2006, 2011; Mohapatra et al., 2016). Here, however, we found that the spread of Na^+^ was largely independent from spine densities, ranging from 3-12 spines/10 µm. A likely reason for this apparent discrepancy is that spine numbers were lower than those in the foresaid studies, which simulated densities >20 spines/10 µm (Santamaria et al., 2011; Mohapatra et al., 2016). However, while overall spine density in the ranges measured apparently does not quantitatively alter bulk diffusion along dendrites, individual spine necks can locally hinder diffusion into the spine head as described above.

Further analysis indicated that the apparent slowing down of Na^+^ diffusion and decrease in *D*_*app*_ were due to the relatively long phase of Na^+^ influx as well as to the restricted lengths of dendrites analyzed. With these boundary conditions taken into account, experimental traces could be fit well assuming a D_Na+_ of 600 µm^2^/s, which corresponds to that reported for muscle cells (Kushmerick and Podolsky, 1969). We therefore conclude that Na^+^ diffusion in dendrites of CA1 pyramidal neurons is well-described by the normal diffusion model. Furthermore, diffusion was independent from the presence of spines at densities found here (≤12 spines/10 µm). Na^+^ diffusion within dendrites, therefore, seems to obey similar biophysical principles as in axons, in which action potential-induced [Na^+^]_i_ transients were replicated assuming lateral diffusion at a D_Na+_ of 600 µm^2^/s (Kole et al., 2008; Fleidervish et al., 2010; Baranauskas et al., 2013; Filipis and Canepari, 2021; Zang and Marder, 2021; Kotler et al., 2023). This is much more rapid as compared to Ca^2+^, for which a cytosolic diffusion coefficient of 250 µm^2^/s was estimated (Allbritton et al., 1992). In contrast to Na^+^, however, intracellular Ca^2+^ movement is restricted owing to immobile buffers (Sabatini et al., 2002).

Na^+^ diffusion was also independent from the average dendrite diameter (1.2-1.8 µm). At this diameter, and a D_Na+_ of 600 µm^2^/s, Na^+^ will be very quickly (less than ~2 ms) equalized across the dendritic radius, which cannot be resolved at the given experimental temporal and spatial resolution. This conclusion is in agreement with reports showing that molecular diffusion was independent on the diameter of dendrites of cerebellar Purkinje cells at diameters of 1.5 - 3.5 µm (Santamaria et al., 2006).

### Relevance of dendritic Na^+^ diffusion

Our simulations confirmed and extended our experimental data by showing that lateral diffusion of Na^+^ is the main mechanism for clearance of local [Na^+^]_i_ increases in early phases after influx, while NKA-mediated transport is more relevant at later stages. Computational modelling also proposed a similar “division of labor” between early diffusion- and late transporter-mediated clearance mechanisms for handling of GABA_A_ receptor-induced Cl^-^ influx (Doyon et al., 2011). Rapid diffusion of Na^+^ will protect synapses from saturation upon repeated activity, i. e., from a reduction in EPSP amplitudes owing to a reduced Na^+^ Nernst potential (Bush and Sejnowski, 1994; Zylbertal et al., 2017).

Fast diffusive clearance of Na^+^ will also dampen Na^+^-dependent stimulation of the NKA, and thereby reduce local ATP consumption (Mondragao et al., 2016; Gerkau et al., 2019). This differs from global Na^+^ signals which result in well-detectable decreases in neuronal ATP (Mondragao et al., 2016; Gerkau et al., 2019; Lerchundi et al., 2020). The data moreover shows that after erosion of concentration gradients, NKA-mediated Na^+^ export becomes more important. This export, will be distributed in time and space, thereby reducing the immediate local metabolic burden.

A reduced net diffusion of Na^+^, either because of spatial constraints or minor concentration gradients, will cause prolonged Na^+^ accumulation and activation of NKA. The hyperpolarization resulting from the latter has been proposed to modulate neuronal activity and to serve as a Ca^2+^-independent form of short-term memory (Pulver and Griffith, 2010; Forrest et al., 2012; Gulledge et al., 2013; Picton et al., 2017; Zylbertal et al., 2017). At the same time, elevation of [Na^+^]_i_ may contribute to intracellular Ca^2+^-signaling due to reversal of Na^+^/Ca^2+^-exchangers (Scheuss et al., 2006; Khananshvili, 2014; Zylbertal et al., 2015; Zylbertal et al., 2017). For a comprehensive understanding of the role of Na^+^ diffusion and NKA activation in shaping Na^+^ transients in dendrites at different forms of activity, however, clearly more experimental and computational studies are required.

## Acknowledgements

This study was supported by the German Research Foundation (DFG), Projekt# 461542557 and Research Unit/FOR 2795 “Synapses under Stress” (Ro2327/13-1, 13-2 to C.R.R. and a Mercator fellowship to G.U.); by the Federal Ministry of Education and Research (BMBF), Germany (Project SynGluCross to C.R.R.); by the National Institutes of Health (R01NS130916 to G.U. and R01NS130759 to F.S.) and by the National Science Foundation (2318139 to F.S.). The authors wish to thank Louis A. Neu and Nils Pape for their help with the preparation of organotypic slice cultures and Claudia Roderigo and Simone Durry for expert technical assistance.

## References

Allbritton NL, Meyer T, Stryer L (1992) Range of messenger action of calcium ion and inositol 1,4,5-trisphosphate. Science 258:1812–1815.

Azarias G, Kruusmagi M, Connor S, Akkuratov EE, Liu XL, Lyons D, Brismar H, Broberger C, Aperia A (2013) A specific and essential role for Na,K-ATPase alpha3 in neurons co-expressing alpha1 and alpha3. J Biol Chem 288:2734–2743.

Baranauskas G, David Y, Fleidervish IA (2013) Spatial mismatch between the Na^+^ flux and spike initiation in axon initial segment. Proc Natl Acad Sci U S A 110:4051–4056.

Barreto E, Cressman JR (2011) Ion concentration dynamics as a mechanism for neuronal bursting. Journal of biological physics 37:361–373.

Blaustein MP, Lederer WJ (1999) Sodium/calcium exchange: its physiological implications. Physiol Rev 79:763–854.

Bouron A, Reuter H (1996) A role of intracellular Na^+^ in the regulation of synaptic transmission and turnover of the vesicular pool in cultured hippocampal cells. Neuron 17:969–978.

Buchin A, de Frates R, Nandi A, Mann R, Chong P, Ng L, Miller J, Hodge R, Kalmbach B, Bose S, Rutishauser U, McConoughey S, Lein E, Berg J, Sorensen S, Gwinn R, Koch C, Ting J, Anastassiou CA (2022) Multi-modal characterization and simulation of human epileptic circuitry. Cell reports 41:111873.

Bush PC, Sejnowski TJ (1994) Effects of inhibition and dendritic saturation in simulated neocortical pyramidal cells. J Neurophysiol 71:2183–2193.

Canepari M, Ross WN (2024) Spatial and temporal aspects of neuronal calcium and sodium signals measured with low-affinity fluorescent indicators. Pflügers Arch 476:39–48.

Cartailler J, Holcman D (2018) Electrical transient laws in neuronal microdomains based on electro-diffusion. Physical chemistry chemical physics : PCCP 20:21062–21067.

Cressman JR, Jr., Ullah G, Ziburkus J, Schiff SJ, Barreto E (2009) The influence of sodium and potassium dynamics on excitability, seizures, and the stability of persistent states: I. Single neuron dynamics. J Comput Neurosci 26:159–170.

David G, Barrett JN, Barrett EF (1997) Spatiotemporal gradients of intra-axonal [Na^+^] after transection and resealing in lizard peripheral myelinated axons. J Physiol 498 (Pt 2):295–307.

Dayan P, Abbott LF (2001) Theoretical neuroscience: Computational and mathematical modeling In, pp 224–225: MIT Press.

Despa S, Bers DM (2003) Na/K pump current and [Na](i) in rabbit ventricular myocytes: local [Na](i) depletion and Na buffering. Biophys J 84:4157–4166.

Despa S, Steels P, Ameloot M (2000) Fluorescence lifetime microscopy of the sodium indicator sodium-binding benzofuran isophthalate in HeLa cells. Anal Biochem 280:227–241.

Diarra A, Sheldon C, Church J (2001) In situ calibration and [H+] sensitivity of the fluorescent Na+ indicator SBFI. Am J Physiol Cell Physiol 280:C1623–1633.

Donoso P, Mill JG, O’Neill SC, Eisner DA (1992) Fluorescence measurements of cytoplasmic and mitochondrial sodium concentration in rat ventricular myocytes. J Physiol 448:493–509.

Doyon N, Prescott SA, Castonguay A, Godin AG, Kroger H, De Koninck Y (2011) Efficacy of synaptic inhibition depends on multiple, dynamically interacting mechanisms implicated in chloride homeostasis. PLoS Comput Biol 7:e1002149.

Filipis L, Canepari M (2021) Optical measurement of physiological sodium currents in the axon initial segment. J Physiol 599:49–66.

Fleidervish IA, Lasser-Ross N, Gutnick MJ, Ross WN (2010) Na^+^ imaging reveals little difference in action potential-evoked Na^+^ influx between axon and soma. Nat Neurosci 13:852–860.

Forrest MD, Wall MJ, Press DA, Feng J (2012) The sodium-potassium pump controls the intrinsic firing of the cerebellar Purkinje neuron. PLoS One 7:e51169.

Gerkau NJ, Lerchundi R, Nelson JSE, Lantermann M, Meyer J, Hirrlinger J, Rose CR (2019) Relation between activity-induced intracellular sodium transients and ATP dynamics in mouse hippocampal neurons. J Physiol 597:5687–5570.

Gonzalez OC, Timofeev I, Bazhenov M (2024) Role of Ion Concentration Dynamics in Epileptic Seizures. In: Jasper’s Basic Mechanisms of the Epilepsies, 5th Edition (Noebels JL, Avoli M, Rogawski MA, Vezzani A, Delgado-Escueta AV, eds), pp 351–364. New York.

Gulledge AT, Dasari S, Onoue K, Stephens EK, Hasse JM, Avesar D (2013) A sodium-pump-mediated afterhyperpolarization in pyramidal neurons. J Neurosci 33:13025–13041.

Hallermann S, de Kock CP, Stuart GJ, Kole MH (2012) State and location dependence of action potential metabolic cost in cortical pyramidal neurons. Nat Neurosci 15:1007–1014.

Harris JJ, Jolivet R, Attwell D (2012) Synaptic energy use and supply. Neuron 75:762–777.

Holcman D, Yuste R (2015) The new nanophysiology: regulation of ionic flow in neuronal subcompartments. Nat Rev Neurosci 16:685–692.

Jaffe DB, Johnston D, Lasser-Ross N, Lisman JE, Miyakawa H, Ross WN (1992) The spread of Na^+^ spikes determines the pattern of dendritic Ca^2+^ entry into hippocampal neurons. Nature 357:244–246.

Jung DW, Apel LM, Brierley GP (1992) Transmembrane gradients of free Na+ in isolated heart mitochondria estimated using a fluorescent probe. Am J Physiol 262:C1047–1055.

Kelly T, Rose CR (2010) Ammonium influx pathways into astrocytes and neurones of hippocampal slices. J Neurochem 115:1123–1136.

Khananshvili D (2014) Sodium-calcium exchangers (NCX): molecular hallmarks underlying the tissue-specific and systemic functions. Pflugers Arch 466:43–60.

Kole MH, Ilschner SU, Kampa BM, Williams SR, Ruben PC, Stuart GJ (2008) Action potential generation requires a high sodium channel density in the axon initial segment. Nat Neurosci 11:178–186.

Kotler O, Khrapunsky Y, Shvartsman A, Dai H, Plant LD, Goldstein SAN, Fleidervish I (2023) SUMOylation of Na(V)1.2 channels regulates the velocity of backpropagating action potentials in cortical pyramidal neurons. eLife 12:e81463.

Krishnan GP, Bazhenov M (2011) Ionic dynamics mediate spontaneous termination of seizures and postictal depression state. J Neurosci 31:8870–8882.

Kushmerick MJ, Podolsky RJ (1969) Ionic mobility in muscle cells. Science 166:1297–1298.

Lamy CM, Chatton JY (2011) Optical probing of sodium dynamics in neurons and astrocytes. Neuroimage 58:572–578.

Langer J, Rose CR (2009) Synaptically induced sodium signals in hippocampal astrocytes in situ. J Physiol 587:5859–5877.

Langer J, Stephan J, Theis M, Rose CR (2012) Gap junctions mediate intercellular spread of sodium between hippocampal astrocytes in situ. Glia 60:239–252.

Langer J, Gerkau NJ, Derouiche A, Kleinhans C, Moshrefi-Ravasdjani B, Fredrich M, Kafitz KW, Seifert G, Steinhauser C, Rose CR (2017) Rapid sodium signaling couples glutamate uptake to breakdown of ATP in perivascular astrocyte endfeet. Glia 65:293–308.

Lasser-Ross N, Ross WN (1992) Imaging voltage and synaptically activated sodium transients in cerebellar Purkinje cells. Proc Biol Sci 247:35–39.

Lennie P (2003) The cost of cortical computation. Curr Biol 13:493–497.

Lerchundi R, Huang N, Rose CR (2020) Quantitative imaging of changes in astrocytic and neuronal adenosine triphosphate using two different variants of ATeam. Frontiers in cellular neuroscience 14:80, doi:10.3389/fncel.2020.00080.

Lerchundi R, Kafitz KW, Farfers M, Beyer F, Huang N, Rose CR (2019) Imaging of intracellular ATP in organotypic tissue slices of the mouse brain using the FRET-based sensor ATeam1.03^YEMK^. Journal of visualized experiments : JoVE 154: doi: 10.3791/60294.

Lezmy J, Harris JJ, Attwell D (2021) Optimising the energetic cost of the glutamatergic synapse. Neuropharmacology 197:108727.

Lobo VM (1993) Mutual diffusion coefficients in aqueous electrolyte solutions. Pure & Appl Chem 65:2613–2640.

Maravall M, Mainen ZF, Sabatini BL, Svoboda K (2000) Estimating intracellular calcium concentrations and buffering without wavelength ratioing. Biophys J 78:2655–2667.

Meier SD, Kovalchuk Y, Rose CR (2006) Properties of the new fluorescent Na^+^ indicator CoroNa Green: comparison with SBFI and confocal Na^+^ imaging. J Neurosci Methods 155:251–259.

Meyer J, Untiet V, Fahlke C, Gensch T, Rose CR (2019) Quantitative determination of cellular [Na^+^] by fluorescence lifetime imaging with CoroNaGreen. J Gen Physiol 151:1319–1331.

Meyer J, Gerkau NJ, Kafitz KW, Patting M, Jolmes F, Henneberger C, Rose CR (2022) Rapid fluorescence lifetime imaging reveals that TRPV4 channels promote dysregulation of neuronal Na(+) in ischemia. J Neurosci 42:552–566.

Migliore M, Messineo L, Ferrante M (2004) Dendritic Ih selectively blocks temporal summation of unsynchronized distal inputs in CA1 pyramidal neurons. J Comput Neurosci 16:5–13.

Migliore M, Hoffman DA, Magee JC, Johnston D (1999) Role of an A-type K+ conductance in the back-propagation of action potentials in the dendrites of hippocampal pyramidal neurons. J Comput Neurosci 7:5–15.

Miyazaki K, Ross WN (2017) Sodium Dynamics in Pyramidal Neuron Dendritic Spines: Synaptically Evoked Entry Predominantly through AMPA Receptors and Removal by Diffusion. J Neurosci 37:9964–9976.

Miyazaki K, Ross WN (2022) Fast Synaptically Activated Calcium and Sodium Kinetics in Hippocampal Pyramidal Neuron Dendritic Spines. eNeuro 9:10.1523/ENEURO.0396-1522.2022.

Mohapatra N, Tonnesen J, Vlachos A, Kuner T, Deller T, Nagerl UV, Santamaria F, Jedlicka P (2016) Spines slow down dendritic chloride diffusion and affect short-term ionic plasticity of GABAergic inhibition. Scientific reports 6:23196.

Mondragao MA, Schmidt H, Kleinhans C, Langer J, Kafitz KW, Rose CR (2016) Extrusion versus diffusion: mechanisms for recovery from sodium loads in mouse CA1 pyramidal neurons. J Physiol 594:5507–5527.

Moshrefi-Ravasdjani B, Hammel EL, Kafitz KW, Rose CR (2017) Astrocyte sodium signalling and panglial spread of sodium signals in brain white matter. Neurochem Res 42:2505–2518.

Naumann G, Lippmann K, Eilers J (2018) Photophysical properties of Na(+)-indicator dyes suitable for quantitative two-photon fluorescence-lifetime measurements. J Microsc 272:136–144.

Neher E, Augustine GJ (1992) Calcium gradients and buffers in bovine chromaffin cells. J Physiol 450:273–301.

Picton LD, Zhang H, Sillar KT (2017) Sodium pump regulation of locomotor control circuits. J Neurophysiol 118:1070–1081.

Pulver SR, Griffith LC (2010) Spike integration and cellular memory in a rhythmic network from Na+/K+ pump current dynamics. Nat Neurosci 13:53–59.

Pusch M, Neher E (1988) Rates of diffusional exchange between small cells and a measuring patch pipette. Pflugers Arch 411:204–211.

Qian N, Sejnowski T (1989) An electro-diffusion model for computing membrane potentials and ionic concentrations in branching dendrites, spines and axons. Biological cybernetics 62:1–15.

Regehr WG (1997) Interplay between sodium and calcium dynamics in granule cell presynaptic terminals. Biophys J 73:2476–2488.

Rose CR, Konnerth A (2001) NMDA receptor-mediated Na^+^ signals in spines and dendrites. J Neurosci 21:4207–4214.

Rose CR, Ziemens D, Verkhratsky A (2020) On the special role of NCX in astrocytes: Translating Na(+)-transients into intracellular Ca(2+) signals. Cell Calcium 86:102154.

Rose CR, Kovalchuk Y, Eilers J, Konnerth A (1999) Two-photon Na^+^ imaging in spines and fine dendrites of central neurons. Pflugers Arch 439:201–207.

Sabatini BL, Maravall M, Svoboda K (2001) Ca(2+) signaling in dendritic spines. Curr Opin Neurobiol 11:349–356.

Sabatini BL, Oertner TG, Svoboda K (2002) The life cycle of Ca(2+) ions in dendritic spines. Neuron 33:439–452.

Santamaria F, Wils S, De Schutter E, Augustine GJ (2006) Anomalous diffusion in Purkinje cell dendrites caused by spines. Neuron 52:635–648.

Santamaria F, Wils S, De Schutter E, Augustine GJ (2011) The diffusional properties of dendrites depend on the density of dendritic spines. Eur J Neurosci 34:561–568.

Scheuss V, Yasuda R, Sobczyk A, Svoboda K (2006) Nonlinear [Ca2+] signaling in dendrites and spines caused by activity-dependent depression of Ca2+ extrusion. J Neurosci 26:8183–8194.

Sejnowski TJ, Qian N (1992) Synaptic integration by electrodiffusion in dendritic spines. In: Single Neuron Computation (McKenna T, Davis J, Zornetzer SF, eds), pp 117–139. Boston: Academic Press.

Sheldon C, Cheng YM, Church J (2004) Concurrent measurements of the free cytosolic concentrations of H(+) and Na(+) ions with fluorescent indicators. Pflügers Arch 449:307–318.

Song H, Thompson SM, Blaustein MP (2013) Nanomolar ouabain augments Ca^2+^ signalling in rat hippocampal neurones and glia. J Physiol 591:1671–1689.

Spruston N (2008) Pyramidal neurons: dendritic structure and synaptic integration. Nat Rev Neurosci 9:206–221.

Stoppini L, Buchs PA, Muller D (1991) A simple method for organotypic cultures of nervous tissue. J Neurosci Methods 37:173–182.

Sweadner K (1995) Sodium, potassium-adenosine triphosphatase and its isoforms. In: Neuroglia (Kettenmann H RB, ed), pp 259–272. London, New York: Oxford University Press.

Tecuatl C, Ljungquist B, Ascoli GA (2024) Accelerating the continuous community sharing of digital neuromorphology data. FASEB Bioadv 6:207–221.

Ullah G, Cressman JR, Jr., Barreto E, Schiff SJ (2009) The influence of sodium and potassium dynamics on excitability, seizures, and the stability of persistent states. II. Network and glial dynamics. J Comput Neurosci 26:171–183.

Wei Y, Ullah G, Schiff SJ (2014) Unification of neuronal spikes, seizures, and spreading depression. J Neurosci 34:11733–11743.

Weilinger NL, Wicki-Stordeur LE, Groten CJ, LeDue JM, Kahle KT, MacVicar BA (2022) KCC2 drives chloride microdomain formation in dendritic blebbing. Cell reports 41:111556.

Yuste R, Denk W (1995) Dendritic spines as basic functional units of neuronal integration. Nature 375:682–684.

Yuste R, Majewska A, Holthoff K (2000) From form to function: calcium compartmentalization in dendritic spines. Nat Neurosci 3:653–659.

Zang Y, Marder E (2021) Interactions among diameter, myelination, and the Na/K pump affect axonal resilience to high-frequency spiking. Proc Natl Acad Sci U S A 118:e2105795118.

Zhao H, Carney KE, Falgoust L, Pan JW, Sun D, Zhang Z (2016) Emerging roles of Na(+)/H(+) exchangers in epilepsy and developmental brain disorders. Prog Neurobiol 138-140:19–35.

Zheng K, Bard L, Reynolds JP, King C, Jensen TP, Gourine AV, Rusakov DA (2015) Time-Resolved Imaging Reveals Heterogeneous Landscapes of Nanomolar Ca(2+) in Neurons and Astroglia. Neuron 88:277–288.

Zhou Z, Neher E (1993) Mobile and immobile calcium buffers in bovine adrenal chromaffin cells. J Physiol 469:245–273.

Zhu Y, Li D, Huang H (2020) Activity and Cytosolic Na(+) Regulate Synaptic Vesicle Endocytosis. J Neurosci 40:6112–6120.

Ziemens D, Oschmann F, Gerkau NJ, Rose CR (2019) Heterogeneity of activity-induced sodium transients between astrocytes of the mouse hippocampus and neocortex: Mechanisms and consequences. J Neurosci 39:2620–2634.

Zylbertal A, Yarom Y, Wagner S (2017) The Slow Dynamics of Intracellular Sodium Concentration Increase the Time Window of Neuronal Integration: A Simulation Study. Frontiers in computational neuroscience 11:85.

Zylbertal A, Kahan A, Ben-Shaul Y, Yarom Y, Wagner S (2015) Prolonged Intracellular Na^+^ Dynamics Govern Electrical Activity in Accessory Olfactory Bulb Mitral Cells. PLoS Biol 13:e1002319.

